# Functional specialization of hippocampal somatostatin-expressing interneurons

**DOI:** 10.1101/2023.04.27.538511

**Authors:** Simon Chamberland, Gariel Grant, Robert Machold, Erica R. Nebet, Guoling Tian, Monica Hanani, Klas Kullander, Richard W. Tsien

## Abstract

Hippocampal somatostatin-expressing (*Sst*) GABAergic interneurons (INs) exhibit considerable anatomical and functional heterogeneity. Recent single cell transcriptome analyses have provided a comprehensive *Sst*-IN subtype census, a plausible molecular ground truth of neuronal identity whose links to specific functionality remain incomplete. Here, we designed an approach to identify and access subpopulations of *Sst*-INs based on transcriptomic features. Four mouse models based on single or combinatorial Cre- and Flp- expression differentiated functionally distinct subpopulations of CA1 hippocampal *Sst-*INs that largely tiled the morpho-functional parameter space of the *Sst*-INs superfamily. Notably, the *Sst;;Tac1* intersection revealed a population of bistratified INs that preferentially synapsed onto fast-spiking interneurons (FS-INs) and were both necessary and sufficient to interrupt their firing. In contrast, the *Ndnf;;Nkx2-1* intersection identified a population of oriens lacunosum-moleculare (OLM) INs that predominantly targeted CA1 pyramidal neurons, avoiding FS-INs. Overall, our results provide a framework to translate neuronal transcriptomic identity into discrete functional subtypes that capture the diverse specializations of hippocampal *Sst*-INs.

**Significance statement:** GABAergic interneurons are important regulators of neuronal activity. Recent transcriptome analyses have provided a comprehensive classification of interneuron subtypes, but the connections between molecular identities and specific functions are not yet fully understood. Here, we developed an approach to identify and access subpopulations of interneurons based on features predicted by transcriptomic analysis. Functional investigation in transgenic animals revealed that hippocampal somatostatin-expressing interneurons (*Sst*-INs) can be divided into at least four subfamilies, each with distinct functions. Most importantly, the *Sst;;Tac1* intersection targeted a population of bistratified cells that overwhelmingly targeted fast-spiking interneurons. In contrast, the *Ndnf;;Nkx2-1* intersection revealed a population of oriens lacunosum-moleculare interneurons that selectively targeted CA1 pyramidal cells. Overall, this study reveals that genetically distinct subfamilies of *Sst*-INs form specialized circuits in the hippocampus with differing functional impact.

## Introduction

A conserved feature of cortical circuits is the presence of numerous excitatory neurons whose activity is kept in check and coordinated by heterogeneous populations of GABAergic INs (1–4). IN heterogeneity is reflected in their neurochemical content, electrophysiological properties, anatomy, and connectivity (1, 2, 5). Because varied combinations of these features determine the specific function of each IN subtype, understanding how neuronal circuits process information requires a functional dissection of IN diversity. *Sst*-INs constitute a major fraction of INs in hippocampal area CA1 where they are largely found in stratum oriens and in the alveus (O/A) (1). As an integral part of the feedback inhibitory circuit, they control dendritic integration and pace network activity (6–8). While *Sst*-INs have been functionally studied as a single ensemble ((6, 9, 10) but see (11, 12)), multiple studies provide clues to divisions in their neurochemical, anatomical and electrophysiological properties (13–19). For example, the overall population of *Sst*-INs can target both principal neurons and FS-INs, resulting respectively in inhibition and disinhibition, two mostly opposing network effects (9, 20–23). Whether specific subtypes of *Sst*-INs account for these disparate circuit functions remains unknown.

Recent single cell transcriptomic studies have provided deep insights into neuronal diversity at the molecular level. Transcriptomic heterogeneity is largely aligned with the traditional subdivision of neurons into superfamilies (24–26), including CA1 hippocampal *Sst*-INs, and indicates the existence of multiple subfamilies with distinct molecular profiles (3). While this transcriptomic classification approach allows for the identification of putative *Sst*-IN subtypes, it inherently lacks the ability to directly predict or investigate functional specialization (3). Thus, a key challenge to understanding how molecularly defined SST-IN subtypes regulate brain circuitry is how to identify and experimentally access these populations *in situ*.

Here, we describe a series of genetic approaches that leverage molecular profiling data to distinguish *Sst*-IN subtypes for experimental interrogation. We dissected the diversity of CA1 hippocampal *Sst*-INs by generating 4 lines of transgenic mice that were predicted to target distinct and minimally overlapping *Sst*-INs subpopulations. Our results revealed that the 4 subtypes of *Sst*-INs largely tile the anatomical and electrophysiological features attributed to *Sst*-INs overall, and reduced the intrapopulation variation of most of the parameters sampled. We discovered that *Sst*-IN subtypes are highly specialized in the neurons they target, exemplified by *Sst;;Tac1* bistratified INs that selectively target and interrupt FS-INs to disinhibit the CA1 microcircuit, in contrast to a novel subclass of *Ndnf;;Nkx2-1* OLMs INs that preferentially innervate and inhibit CA1 pyramidal neurons (CA1-PYRs).

## Results

### A genetic dissection of *Sst*-IN diversity

In the CA1 hippocampus, *Sst*-INs adopt multiple anatomical phenotypes defined by their axonal projection (13, 18). Whether anatomical differences can be aligned with genetically distinct neuronal subpopulations within the *Sst*-IN superfamily remains unclear.

To investigate the anatomical diversity of CA1 O/A *Sst*-INs, we bred *Sst*-Cre animals to the Ai9 reporter line and performed whole-cell recordings with biocytin fills from TdTomato+ INs in acute hippocampal slices (Fig. 1A). Post-hoc anatomical tracings confirmed previous reports that hippocampal CA1 *Sst*-INs exhibit diverse axonal projection patterns (n = 25; Fig. 1B and Fig. S1) (13, 14, 27). Anatomical heterogeneity of *Sst*-INs is paralleled at the transcriptomic level, and a large single-cell transcriptomic dataset containing CA1 *Sst*-INs is publicly available (3) (henceforth referred to as the Harris dataset). Transcriptomic datasets allow genetically similar neurons to be put closest to each other in principal component space, in turn represented on plots that render multi-dimensional information on 2D maps. We reasoned that genes or gene pairs that map onto restricted clusters of neurons and minimize intracluster distances might be good predictors of constituent subpopulations of neurons that later prove to be functionally different. First, we used spatial dispersion statistics to uncover genes and pairs of genes that minimized both the standard distance and the inter-quartile distance on the 2D map in the Harris dataset, agnostic of gene identity (Fig. 1C, and Figs. S2, S3). Second, we mapped neurons expressing these genes and visually selected distinct populations (Fig. 1D). Consequently, we identified multiple combination of genes that tiled the general population of *Sst*-INs (Fig. 1D) with minimal overlap at the individual cell level (Fig. 1E). To test the hypothesis that these genetic features identify functionally distinct *Sst*-INs subpopulations, we generated transgenic mice based on combinatorial expression of Cre- and Flp- recombinases (28). We therefore generated *Sst-Flp*;;*Tac1-Cre*, *Ndnf-Flp*;;*Nkx2-1-Cre* and *Sst-Flp;;Nos1-Cre* transgenic lines (referred to as *Sst;;Tac1*, *Ndnf;;Nkx2-1* and *Sst;;Nos1*); we further leveraged the existing *Chrna2-Cre* line, motivated by the observation that *Chrna2* was one of the top ranked genes in our screening and prior knowledge that this transgenic line targets a specific subtype of *Sst*-IN (11).

**Figure 1:**
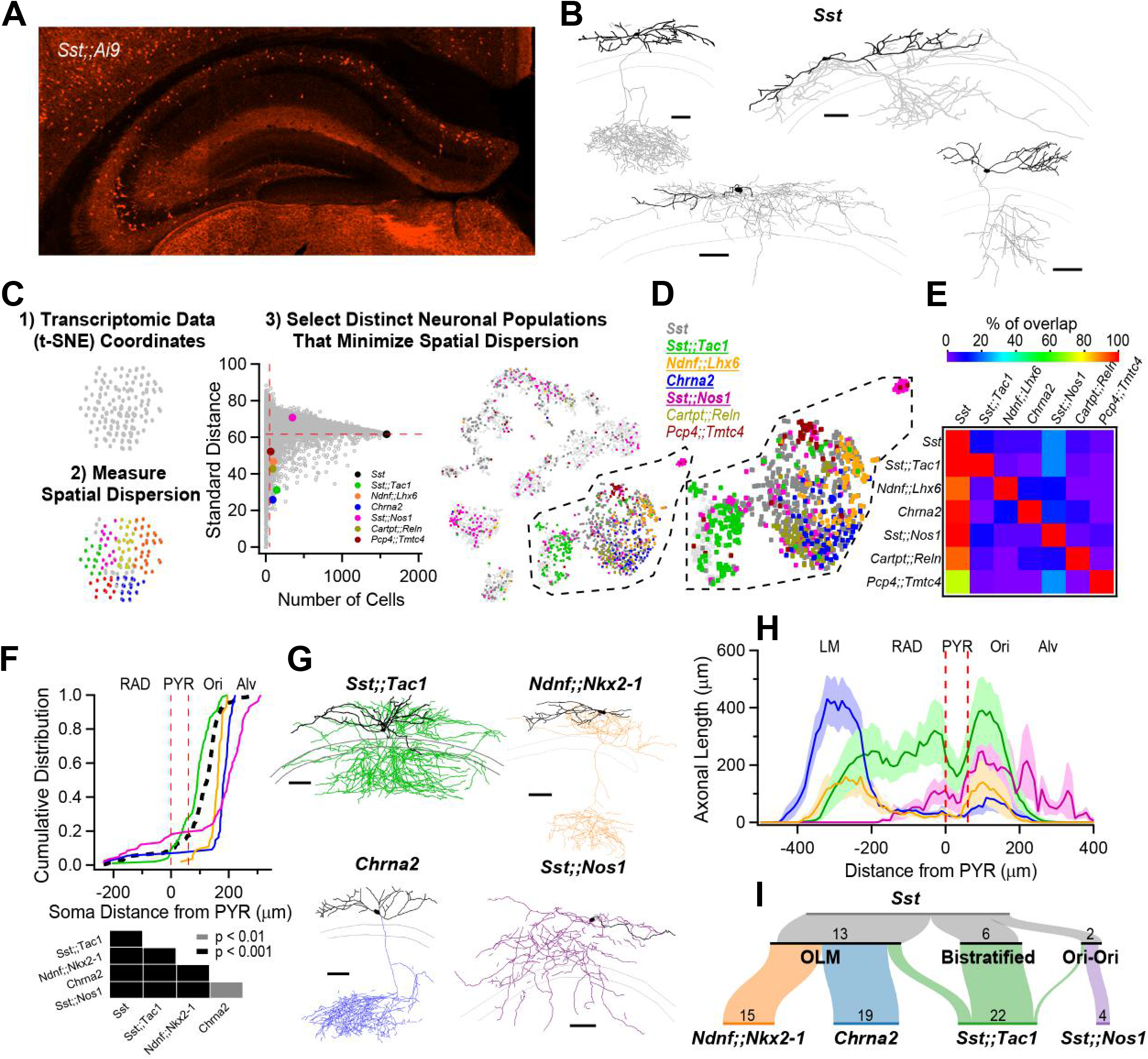
Anatomical heterogeneity of hippocampal *Sst*-INs is partly solved by linking genetic identity to function. **A**, Confocal image from a *Sst;;Ai9* mouse brain microsection showing the distribution of hippocampal neurons expressing the fluorescent protein TdTomato. In the CA1 region, *Sst*-INs are mostly found in stratum oriens/alveus (O/A). **B**, Neurolucida reconstructions of CA1 O/A INs recorded in the *Sst;;Ai9* mouse model and filled with biocytin. Individual examples selected to highlight the diversity of axonal projections from these neurons (dendrites in black, axon in gray). Calibration bars = 100 μm. **C**, Strategy to identify genes or pairs of genes delineating clusters of neurons that tile the larger *Sst*-IN population in the Harris *et al.* dataset (see Methods). **D**, Selection of gene pairs to generate intersectional transgenic mouse models (bold and underlined). The gene *Chrna2* by itself fulfills the established criteria and enabled the use of a pre-existing transgenic mouse line (*11*). **E**, Matrix showing little overlap of subsets of neurons expressing the selected combination of genes. Two potential gene pairs additionally identified within the Harris dataset are shown. Percentage of overlap color coded, where red represents 100% overlap and violet represents 0% overlap. Percentages normalized relative to diagonal (100%). **F**, Quantification of the localization of fluorescently labelled cell bodies in the five genotypes relative to the PYR layer in the CA1 hippocampus. Table below reports the p-values from KS tests between the five genotypes after Holm-Bonferroni correction for multiple comparisons. **G**, Neurolucida reconstructions of representative interneurons visually targeted for recording by the expression of a fluorescent reporter in the different transgenic mouse models. Individual neurons were recorded and filled with biocytin (axons colored according to genotype, dendrites in black). Calibration bars, 100 μm. **H**, Histogram of axonal distribution for all interneurons recorded and filled in the four transgenic mouse models as a function of distance from the pyramidal cell layer (indicate by the dashed red lines). The shaded areas correspond to the standard error. **I**, Sankey diagrams showing the segregation of *Sst*-INs into three broadly defined anatomical categories, OLM, bistratified and oriens-oriens (top); the genetically identified subclasses (bottom) capture and tile the three general anatomical categories of *Sst*-INs, and further refine the within-genotype anatomical identity. The number of recorded and identified neurons is shown.

We bred *Sst;;Tac1*, *Ndnf;;Nkx2-1*, *Sst;;Nos1* and *Chrna2* mice to reporter lines (Ai65 for dual Cre-/Flp- recombinases and Ai9 for single Cre- recombinase), resulting in the expression of TdTomato in these neurons. Measuring the location of TdTomato+ INs as a function of distance from the pyramidal cell layer showed a cell type-specific distribution that largely tiled the general *Sst*-IN population (Fig. 1F). While *Sst;;Tac1*-INs were located closer to the CA1 pyramidal layer, *Ndnf;;Nkx2-1*-INs and *Chrna2*-INs were found progressively deeper in O/A; in contrast, *Sst;;Nos1*-INs were found mostly in the alveus, with some neurons sparsely distributed in strata radiatum and lacunosum-moleculare (LM) (Fig. 1F).

We next investigated the anatomy of neurons identified in transgenic lines with whole-cell recordings and biocytin fills, focusing on cells bodies within O/A, followed by post-hoc anatomical reconstruction. In all cases, the axonal distribution revealed a preference for dendritic layers (Fig. 1G, Figs. S4-7), a feature typical of *Sst*-INs. Quantifying the axonal distribution across the CA1 layers revealed four distinct axonal projection patterns: 1) *Sst;;Tac1*-INs overwhelmingly targeted strata oriens and radiatum; 2) *Ndnf;;Nkx2-1*-INs projected axons to both strata oriens and LM; 3) *Chrna2*-INs exhibited a strong and almost exclusive axonal projection to LM; and 4) *Sst;;Nos1*- INs mostly innervated stratum oriens (Fig. 1H, Fig. S8, Supplementary Table 1). Finally, we associated the genetic identities of INs with commonly used anatomical nomenclature. The *Sst*- INs superfamily contained neurons from the OLM (n = 14), bistratified (n = 8) and oriens-oriens (n = 3) subtypes (Fig. 1I, top). We found that the OLMs were constituted by *Ndnf;;Nkx2-1*-INs (n = 15/15) and *Chrna2*-INs (n = 19/19), while the bistratified and oriens-oriens categories were disproportionately and almost exclusively represented by *Sst;;Tac1*-INs (n = 18/23) and *Sst;;Nos1*-INs (n = 4/4), respectively (Fig. 1I, bottom). Therefore, the wide-ranging anatomical features of *Sst*-INs can be accounted for by the more narrowly defined morphologies of the genetically defined subtypes.

### Electrophysiological features of Sst-INs subpopulations explain the observed variation within the superfamily

*Sst*-INs are generally known as regular-spiking INs and demonstrate a large hyperpolarization-activated cation current (I_h_) (29). Variations in the firing patterns of *Sst*-INs have been reported before (17) and likely contribute to cell type-specific recruitment of these neurons during hippocampal activity (15, 30, 31), but whether the variation within the superfamily can be attributed to genetically defined cells remains unknown.

We next investigated the electrophysiological profiles of *Sst*-INs subtypes and compared them to the superfamily (Fig. 2A). While the firing frequency increased similarly with current injection across all *Sst*-INs subtypes (Fig. 2B), *Sst;;Nos1* demonstrated marked depolarization block (Fig. 2B). We next analyzed typical action potential (AP) parameters and compared their intrapopulation variance (Fig. 2C-F and Fig. S9). Cell type-specific differences were evident (Fig. 2D-F, Fig. S9, Supplementary Table 2). For example, the AP maximal rate of fall was significantly different between all subpopulations (Fig. 2D; KS test: p < 0.05; statistical treatment of complete data set in Supplementary Table 2). In addition, the collective electrophysiological properties of these neurons largely accounted for the range of parameters found in the *Sst*-INs superfamily overall (Fig. 2D-F and Fig. S9). Furthermore, the coefficient of variation (CV) for these parameters was generally lower for all *Sst*-IN subpopulations (Fig. 2D-F and Fig. S9) compared to the superfamily (in 27 out of 32 cases). The tiling was sometimes incomplete (Fig. 2E), aligning with the fact that the four transgenic lines only partly cover the full transcriptomic space of the *Sst*-IN superfamily (Fig. 1D,E). Overall, our recordings uncovered cell type-specific differences between *Sst*-IN subpopulations that help explain the variation of electrophysiological parameters within the superfamily.

**Figure 2:**
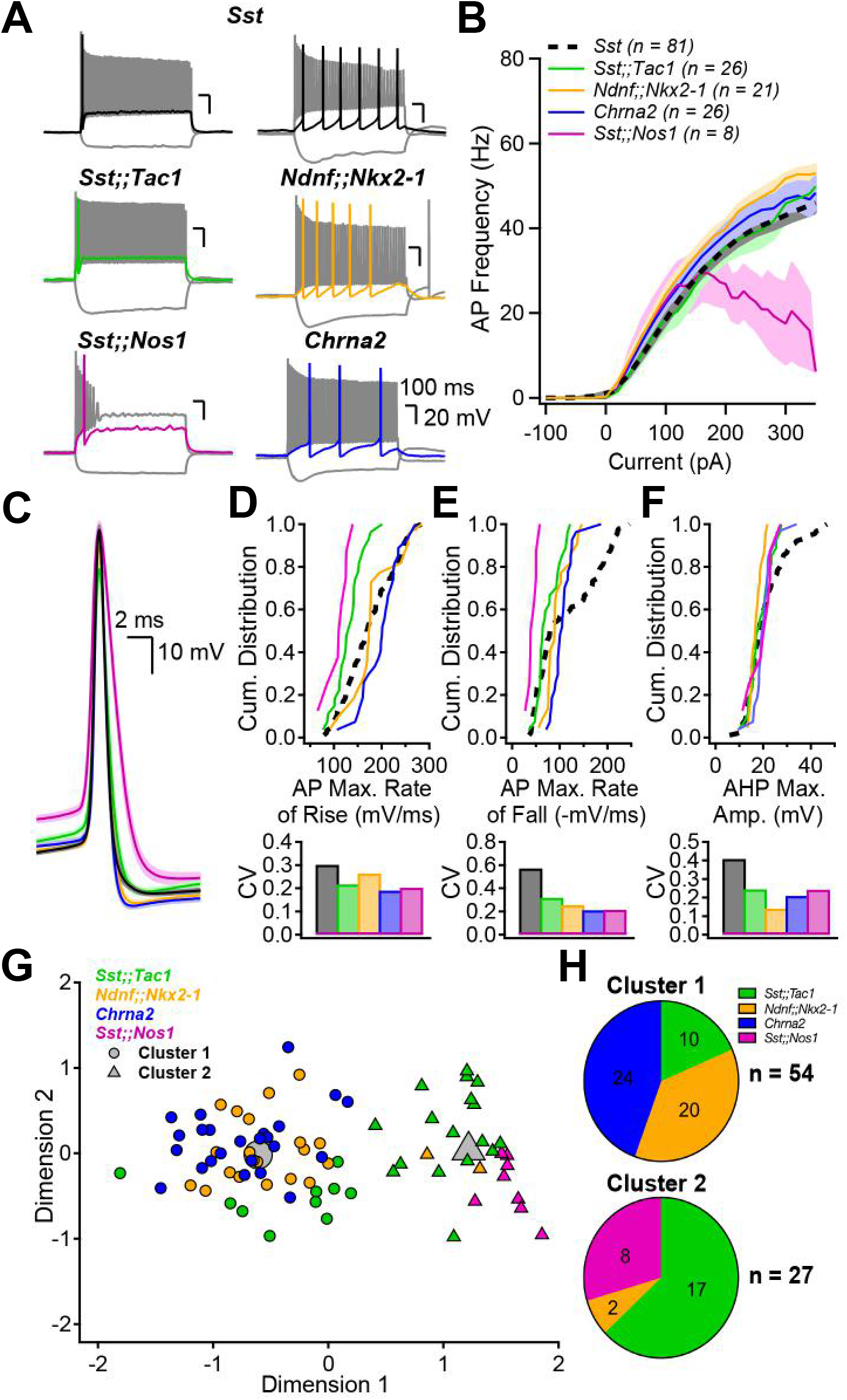
Genetically defined subpopulations of *Sst*-INs tile the electrophysiological parameter space and account for the heterogeneity within the *Sst* family. **A,** Membrane potential changes resulting from hyperpolarizing and depolarizing current pulses in the five transgenic mouse models. Each panel includes a response to hyperpolarizing pulse driving V_m_ between −100 and −90 mV, a response to rheobase current pulse (color), and the maximal firing rate response (gray). **B**, Firing frequency as a function of current injection amplitude. Number of averaged cells is shown. **C**, Action potential waveforms elicited by rheobase current, aligned at peak overshoot, averaged across all interneurons in each subgroup. Shaded areas correspond to standard error. **D**, top, Cumulative distribution of the AP maximal rate of rise (mV/ms) for the five genotypes and associated coefficients of variation (CV, bottom). **E**, **F**, Same as D but for the AP maximal rate of fall (E) and the AP afterhyperpolarization maximal amplitude (F). **G**, Principal component analysis followed by unsupervised k-means clustering analysis using the electrophysiological parameters above and in Figure S9 divides the neurons into two clusters. **H**, Pie charts summarizing the distribution of the genetically identified interneuron subgroups across electrophysiologically-defined clusters. The distribution of neurons was significantly different than expected by chance (Chi-square = 44.485, p < 0.001)

We performed an unsupervised k-means cluster analysis to objectively assign the recorded neurons to groups and probe how much *Sst*-IN subpopulations could be distinguished on the basis of electrophysiological parameters alone (Fig. 2G). First, principal component analysis was performed on the eight electrophysiological parameters measured (Supplementary Table 3). K-means clustering using the first four principal components, which captured more than 90% of the variance, suggested the existence of two distinct clusters (elbow method). Cluster 1 incorporated all the *Chrna2*-INs (24/24) and almost all *Ndnf;;Nkx2-1*-INs (20/22). Cluster 2 captured all the *Sst;;Nos1*-INs (8/8) and most, but not all *Sst;;Tac1*-INs (17/27), far from random overall (p<0.00001 by c^2^ test). Thus, unbiased k-means cluster analysis indicated that our genetically based sorting of *Sst*-IN subpopulations aligned in large part with segregation solely based on electrophysiological properties.

### Cell type-specific targeting by subpopulations of *Sst*-INs

We and others have previously shown that the superfamily of *Sst*-INs targets both CA1-PYRs and FS-INs in the CA1 region (9, 20). In our recent study (20), a small dataset of paired-recordings suggested that *Sst*-expressing bistratified but not OLM cells targeted FS-INs, hinting at cell type-specific connectivity. It remains unknown whether *Sst*-IN subtypes generally provide non-selective or cell type-specific inhibition to their targets.

Optogenetic circuit mapping revealed clear target preference amongst *Sst*-INs subfamilies (Fig. 3). Postsynaptic targets were visually identified and electrophysiologically confirmed as CA1 pyramidal cells (CA1-PYRs), FS-INs and RS-INs with a hyperpolarizing sag (putative *Sst*-INs) before performing voltage-clamp recordings at 0 mV. Optogenetic stimulation (20 ms) of presynaptic *Sst;;Tac1*-INs revealed large amplitude inhibitory postsynaptic currents (IPSCs) in FS-INs (116.1 ± 27.7 pA, n = 20), yet with the same photostimulation, significantly smaller IPSCs in CA1-PYRs (20.8 ± 6.4 pA; n = 12; p < 0.001, Mann Whitney U test) and RS-INs (12.2 ± 2.8 pA; n = 24; p < 0.001, Mann Whitney U test; Fig. 3A-B). In sharp contrast, photostimulation of *Ndnf;;Nkx2-1*-INs generated significantly larger IPSCs in CA1-PYRs (21.7 ± 2.8 pA; n = 14) than in FS-INs (10.2 ± 2.3 pA; n = 9; p < 0.01, Student’s t-test) or RS-INs (1.6 ± 0.9 pA; n = 5; p < 0.001, Mann Whitney U test; Fig. 3A-B). On the other hand, optogenetic stimulation of *Chrna2*- INs resulted in similar IPSCs in CA1-PYRs (25 ± 5.6 pA; n = 10) and FS-INs (21.6 ± 3.9 pA; n = 12; p > 0.4, Mann Whitney U test), that were both much larger than the IPSCs recorded in RS- INs (0.9 ± 0.7 pA; n = 3; p < 0.05 vs. CA1-PYRs and p < 0.01 vs. FS-INs, Mann Whitney U test). Finally, photostimulation of *Sst;;Nos1*-INs revealed almost undetectable IPSCs in the three targets (CA1-PYRs: 0.6 ± 0.5 pA; n = 4; FS-INs: 0.5 ± 0.4 pA, n = 15; RS-INs: 0 pA, n = 9) despite obvious axonal arborization in O/A. To ask how well the subtypes accounted for the impact of *Sst*- positive neurons as a whole, we calculated the sum of IPSC amplitudes evoked by *Sst;;Tac1*-INs, *Ndnf;;Nkx2-1*-INs, *Chrna2*-INs and *Sst;;Nos1*-INs (Fig.3B, red dotted lines labeled S). The summed subgroup events represented 85% of the IPSC in CA1-PYRs directly recorded upon by optogenetic stimulation of the general Sst-IN population; for FS-INs the corresponding percentage was 75%. This suggests that our strategy captured the bulk of *Sst*-INs innervating CA1-PYRs and FS-INs. Moreover, the four *Sst*-INs subtypes hardly influenced regular-spiking INs with a hyperpolarizing sag (Fig. 3A-B), consistent with the idea that *Sst*-INs mostly avoid synapsing with each other (32).

**Figure 3:**
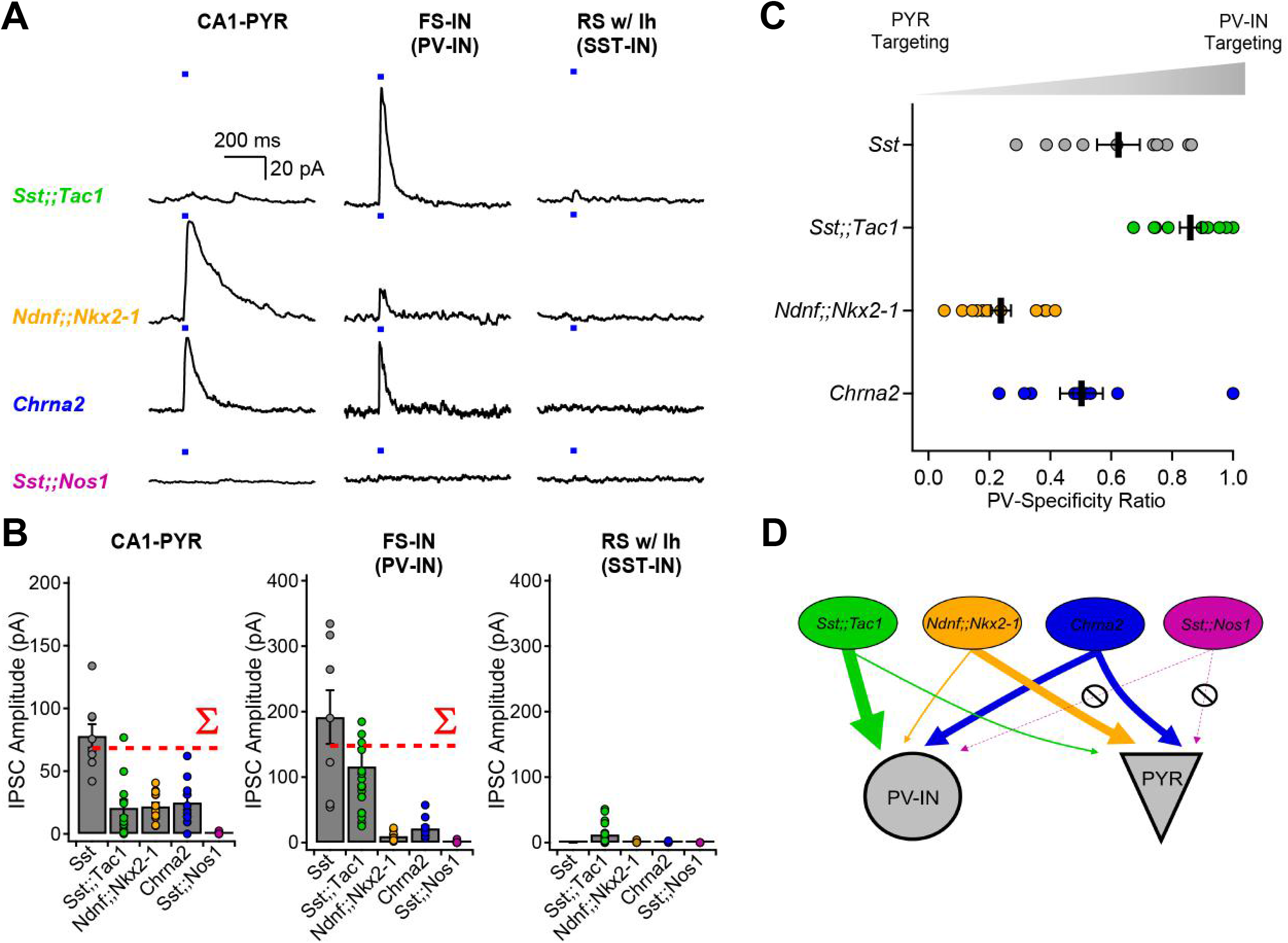
Optogenetic circuit mapping reveals that postsynaptic targets of Sst-INs are subpopulation-specific. **A**, Voltage-clamp recordings (holding potential, 0 mV) from pyramidal cells, fast-spiking interneurons and regular-spiking interneurons with prominent sag, showing representative IPSCs generated by optogenetic activation of IN subpopulations. **B**, Summary bar graph of IPSC amplitudes recorded in the three target types. The dotted red lines show the arithmetic sums of IPSCs generated by photostimulation of the individual subpopulations. **C**, Sequential recordings from neighboring fast-spiking interneurons and pyramidal cells reveals target-specificity of *Sst;;Tac1*-INs, *Ndnf;;Nkx2-1*-INs and *Chrna2*-INs. **D**, Cartoon depicting the target selectivity of *Sst*-IN subpopulations.

For a direct comparison of the relative preference for FS-INs and PYRs, we performed sequential recordings of IPSCs from neighboring CA1-PYRs and FS-INs in response to identical optogenetic stimulation. We analyzed the synaptic strength in these pairs by determining the ratio (IPSC_FS-IN_ / (IPSC_FS-IN_ + IPSC_PYR_)) as an index of FS-IN preference, 0.5 indicating no preference. This normalization circumvented potential confounds including different transgenic animal models, number of presynaptic axons in the slice and optrode placement (Fig. 3C). These experiments confirmed a strong preference of *Sst;;Tac1*-INs for FS-INs over CA1-PYRs (ratio of 0.86 ± 0.3; n = 10 pairs). In contrast, *Ndnf;;Nkx2-1*-INs were found to preferentially target CA1- PYRs (ratio of 0.24 ± 0.03; n = 12 pairs; p < 0.001; Fig. 3C-D), while *Chrna2*-INs contacted both FS-INs and CA1 pyramidal cells without clear preference (ratio of 0.50 ± 0.07; n = 9; Fig. 3C-D). These results, obtained with optogenetic stimulation held fixed, provide strong evidence that *Sst*- IN subpopulations vary widely in the degree to which they target other neuron types and are thus functionally specialized.

### *Sst;;Tac1*-INs are necessary and sufficient to interrupt FS-INs

We now turn to the use of subgroup-specific mouse lines as experimental tools. We recently reported that FS-INs undergo a strikingly persistent interruption of firing upon brief synaptic inhibition, resulting in CA1-PYR disinhibition (20). The interruption of firing was induced by optogenetic stimulation of the general *Sst*-INs population, but whether this function is exclusive or shared amongst multiple *Sst*-INs subpopulation remains unclear.

FS-INs were depolarized to trigger their characteristic fast-spiking and non-adapting firing patterns, and presynaptic subpopulations of *Sst*-INs were optogenetically stimulated. We found that photostimulation of subgroups failed to induce the interruption of firing in the case of *Ndnf;;Nkx2-1*-INs (0% likelihood, n = 5), *Chrna2*-INs (0.9 ± 0.8% likelihood, n = 11) and *Sst;;Nos1*- INs (0% likelihood, n = 4) (Fig. 4A-C). In contrast, *Sst;;Tac1*-INs reliably generated the interruption of firing (77 ± 7% likelihood, n = 15; Fig. 4A-C). Thus, *Sst;;Tac1*-INs triggered the interruption of firing with similar likelihood and dynamics (Fig. 4B,C) as the general *Sst*-INs population (86.1% ± 2.4%, n = 29, p > 0.1) (*Sst* data previously reported in ref. 18). We conclude that among the *Sst*- INs subgroups, the *Sst;;Tac1*-IN subgroup was specifically necessary (Fig. 4A,C) and quantitatively sufficient (Fig. 4B,C) to reliably trigger the persistent interruption of firing. Therefore, these results establish *Sst;;Tac1*-INs in the CA1 hippocampus as a novel subclass of disinhibitory interneurons, one imbued with a potent capability to relieve pyramidal neurons from inhibition (20).

**Figure 4:**
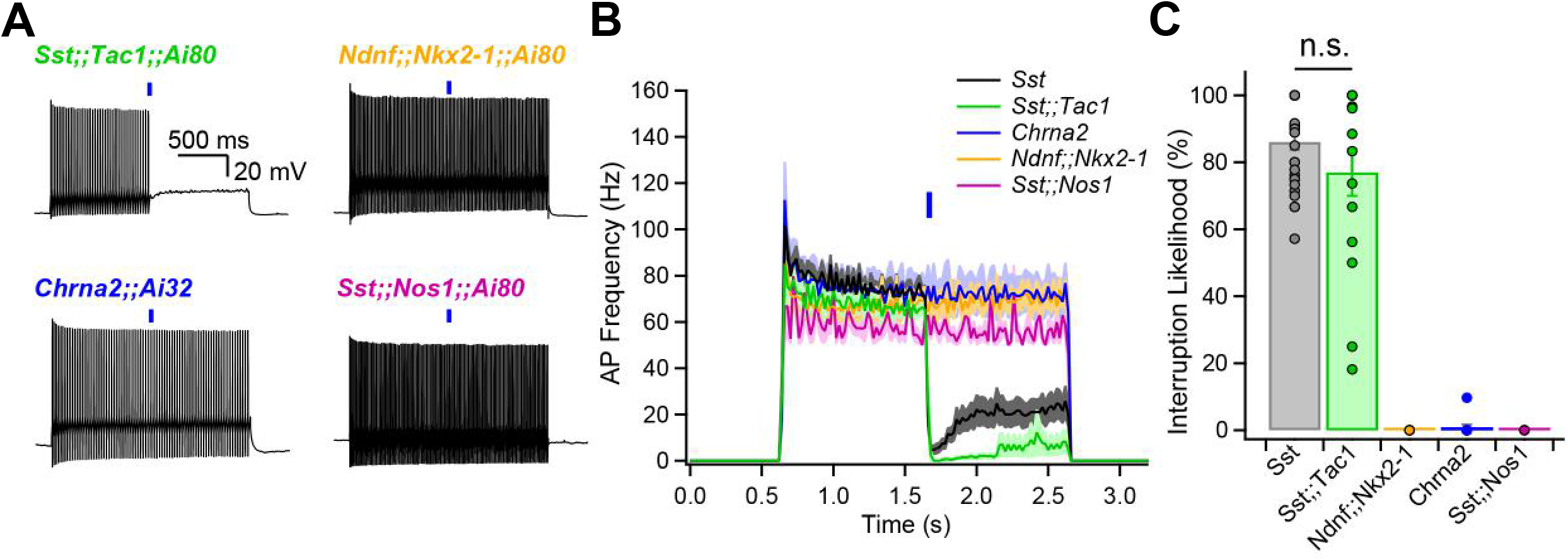
*Sst;;Tac1*-INs are sufficient to interrupt fast-spiking interneurons. **A**, Representative examples showing optogenetic activation of *Sst*-INs subpopulations during sustained fast-spiking interneuron firing evoked by steady current. **B**, Histogram showing the average firing as a function of time before and after optogenetic stimulation of *Sst* subpopulations. **C**, Summary bar graph indicating that *Sst;;Tac1*-INs are sufficient within the general *Sst*-INs population to interrupt fast-spiking interneurons.

## Discussion

Vast heterogeneity amongst hippocampal INs has been identified based on anatomical, neurochemical, electrophysiological and functional criteria (1, 2, 33). Single cell transcriptomic analysis provided a likely complete survey of these cells (3), on which we performed spatial dispersion statistics to predict genetic features identifying minimally overlapping *Sst*-INs subpopulations. To test the hypothesis that these genetic features provide labels to access functionally distinct *Sst*-INs subpopulations, we generated and leveraged transgenic animals. The mouse lines we assembled largely recapitulate *Sst*-INs’ overall synaptic weight and broad spectrum of features: the four tagged subpopulations are distinguishable by a combination of cell-autonomous features, output connectivity and functional impact (Fig. 5). In brief, the *Sst;;Tac1* line labeled bistratified INs, the first genetically-driven access to a population of bistratified neurons. The *Sst;;Nos1* line tagged INs with somata closest to the alveus and diffuse axonal trees, easily told apart from other *Sst*-INs. Two other subtypes shared OLM morphology but were readily distinguished based on their target specificity: *Ndnf;;Nkx2-*1-INs preferentially targeted PYRs over FS-INs, while *Chrna2*-INs (11) lacked PYR:FS-IN preference.

**Figure 5:**
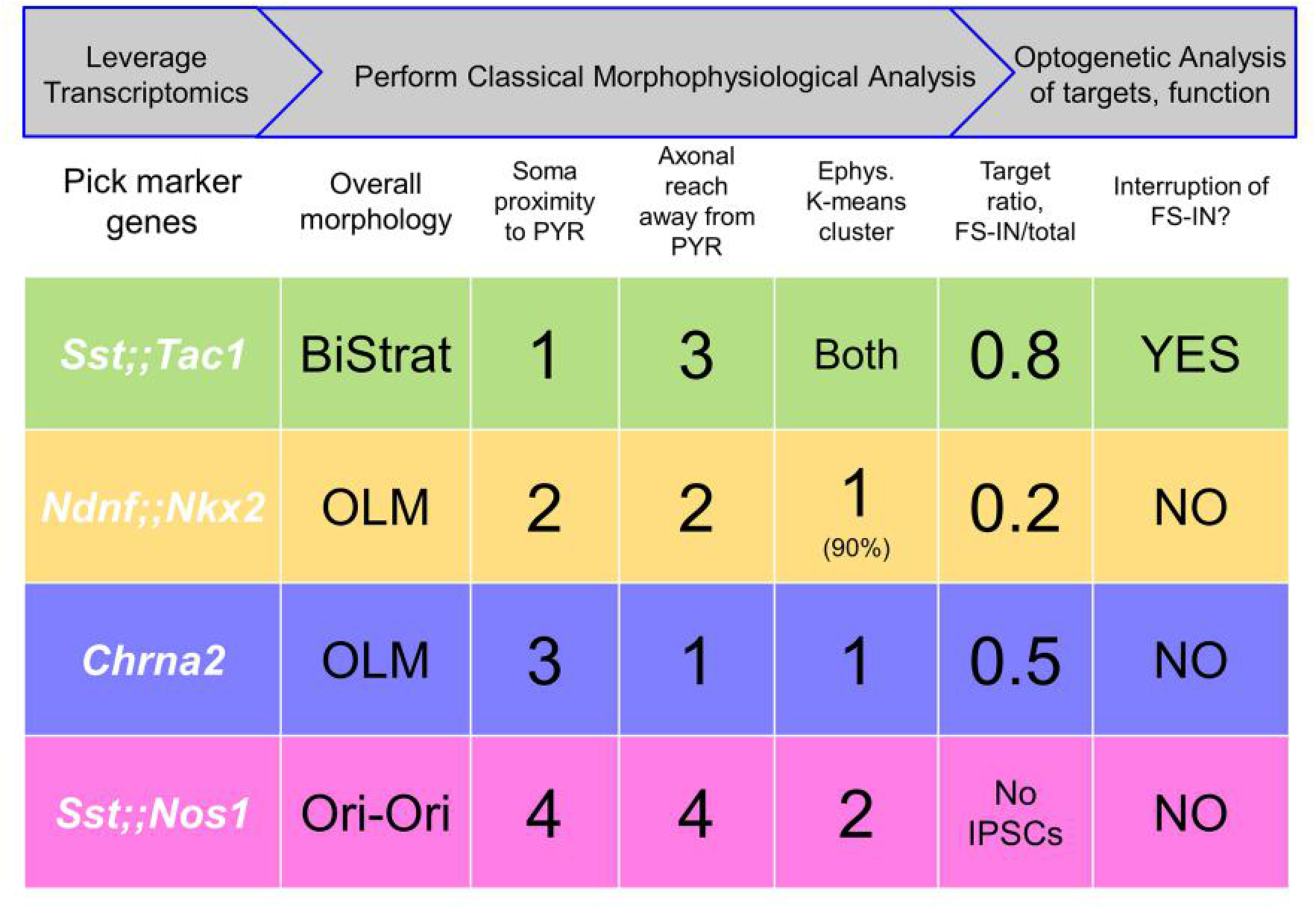
Approach to subdivide a neuronal family into functionally distinct subclasses based on transcriptomics, morphophysiological analysis and optogenetic assessment of impact. **Top rows**, Summary of overall workflow (gray arrows) and operational steps. **Bottom rows**, Summary of 4 subpopulations and some defining characteristics, including morphological ranking with regard to soma proximity to pyramidal layer and axonal extension away from pyramidal layer (Fig. 1); membership in electrophysiological clusters (Fig. 2); optogenetically assessed postsynaptic targeting (Fig. 3); and functional impact (Fig. 4).

Bistratified *Sst;;Tac1* neurons largely spared CA1-PYRs, but preferentially targeted FS *Pv*-INs. This suggests that *Sst;;Tac1* bistratified cells are distinct from *Pv* bistratified cells (34, 35). Thus, Sst;;Tac1-INs are particularly well-suited for disinhibition of CA1 PYRs (20), like subpopulations of *Vip*-INs (36–39). These two types of INs might play complementary circuit roles: *Sst;;Tac1*-INs prefer FS *Pv*-INs over RS *Sst*-INs (Fig. 3C – D), converse to disinhibitory *Vip*-INs, which preferentially innervate RS *Sst*-INs over FS *Pv*-INs (20). Having intersectional mouse lines ready for optogenetic or pharmacogenetic manipulation will hasten future testing of such circuit predictions and their behavioral implications.

Together, the *Ndnf;;Nkx2-1*-OLMs and *Chrna2*-OLMs divide the OLM subtype into two functionally distinct populations (Fig. 5). With the benefit of large numbers of genetically marked cells, we found significant differences in somatic location (Fig. 1F), axonal apportionment (Fig. 1G), electrophysiological properties (Fig. 2) and target specificity (Fig. 3). It is interesting to compare these findings with studies that start with morphofunctionally identified OLM INs (40), or that emphasize developmental origin or expression of ionotropic 5HT3aR serotonin receptors (16). Knowing the genetic profile of *Ndnf;;Nkx2-1*-OLMs and *Chrna2*-OLMs (3) provides a potential starting point but neither of these subpopulations show a pattern of 5HT3aR transcript expression or of origin from the caudal ganglionic eminence. Examination of other tiles in the mosaic of *Sst*-INs (Fig. 1), alert for additional OLM subtypes, would be a logical next step before drawing firm conclusions.

Our findings show practical outcomes of a strategy that leverages single cell transcriptomics (3), classical morpho-physiological analysis (1, 2, 5), and functional connectivity of neuronal subgroups (workflow in first row of Fig. 5). When we began, there was no *a priori* guarantee that tiling based on genetic markers would generate subgroups set apart by morpho-physiological distinctions, as our experiments showed. We suspect that the success of this strategy was not fortuitous--marker genes may reflect deeper differences in gene expression, extending to mechanistically important genes for ion channels, adhesion proteins and developmentally critical transcription factors, etc. (41, 42). A fully bottom-up approach might seem less chancy, but knowledge of genotype-phenotype relationships is still too primitive to support this route. Meanwhile, there may be merit in the pragmatic strategy of using transcriptomic data to predict genetic features identifying distinct and minimally overlapping *Sst*-INs subpopulations and taking the calculated risk of generating intersectional transgenic animals. The animal lines are themselves an end product amenable to functional analysis, both by classic single cell approaches, and by optogenetics on pooled subgroups to determine output connectivity and functional impact (Figs. 3, 4). Like any iterative process of divide and conquer (e.g. Twenty Questions or expression cloning), the assignment of functional roles to ever narrower subgroups might be achieved via multiple paths even if the end result is unique. Having a functional assay (e.g. Fig. 4) provides empirical guidance for the winnowing down procedure and guards against oversplitting (43).

The study of interneuronal function has been greatly accelerated by the development of transgenic animals coupled with optogenetics (44–48), enabling *in situ* identification and selective manipulation of sparse neuronal types (2, 25, 49). Our findings revealed that the transcriptomic profiles of neurons have predictive value for accessing and characterizing subpopulations of neurons, gained via transgenic animals or potentially other approaches (50–53). The tiling strategy developed here to dissect *Sst*-INs could be extended to other groups of neurons, in other brain regions. Genetic access to functionally unified groups of neurons will expedite dissection of circuit function and clarify overriding relationships between neuronal structure and function (11, 12, 54, 55).

## Material and Methods

### Animals and breeding strategies

All experiments performed here were approved performed by the Institutional Animal Care and Use Committee (IACUC) at New York University Langone Medical Center. The experiments reported in this paper involved the use of 11 transgenic mouse lines. *Sst;;Tac1* animals were obtained by crossing *Sst*-Flp (Sst^tm3.1(flpo)Zjh^/AreckJ, JAX stock #28579, (28)) with *Tac1*-Cre (B6;129S-Tac1^tm1.1(cre)Hze^/J, JAX #021877, (56)) mice, and were maintained as double homozygous. *Ndnf;;Nkx2-1* animals were obtained by crossing *Ndnf*-Flp with *Nkx2-1*-Cre (C57BL/6J-Tg(Nkx2-1-cre)2Sand/J, JAX# #008661, (57)) animals. *Ndnf*-Flp animals were generated in collaboration with the New York University Langone Medical Center Rodent Genetic Engineering Laboratory. In brief, a T2A-Flpo-pA cassette was inserted immediately following the last codon in the NDNF open reading frame via homologous recombination in ES cells (B4), followed by clone selection and germline transmission from chimeric founders to establish the colony. *Sst;;Nos1* animals were obtained by crossing *Sst*-Flp to *Nos1*-CreER (B6;129S- Nos1^tm1.1(cre/ERT2)Zjh^/J, JAX stock #014541, (47)) animals. *Sst;;Nos1* animals were maintained as homozygous for *Sst*-Flp and heterozygous for *Nos1*-CreER; double homozygous animals were not viable in our initial observations. *Chrna2*-Cre (Tg(Chrna2-cre)1Kldr) were generated in Uppsala University (Sweden) (11) and maintained as hemizygous. These animals were then bred to the following homozygous reporter lines: Ai9 (B6.Cg-Gt(ROSA)26Sor^tm9(CAG-tdTomato)Hze^/J, JAX stock #007909, (58)), Ai65 (B6;129S-Gt(ROSA)26Sor^tm65.1(CAG-tdTomato)Hze^/J, JAX stock #021875, (59)), Ai32 (B6.Cg-Gt(ROSA)26Sor^tm32(CAG-COP4*H134R/EYFP)Hze^/J, JAX stock # 024109, (60)), Ai80 (B6.Cg-Gt(ROSA)26Sor^tm80.1(CAG-COP4*L132C/EYFP)Hze^/J, JAX stock #025109, (46)). Tamoxifen was administered to *Sst;;Nos1* animals to induce recombination. Tamoxifen (Sigma, TK) was diluted in corn oil at 20 mg/ml, in a heated (55°C) water bath by vortexing every two hours. Animals were gavaged every other day with three doses of 0.15 ml tamoxifen-containing corn oil. P20-35 animals were used for experiments described below.

### Acute hippocampal slice preparation

For acute slice preparation, animals were deeply anesthetized with isoflurane before decapitation. The brain was rapidly extracted into a sucrose-based ice-cold and oxygenated (95% O2, 5%CO2) artificial cerebrospinal fluid (sucrose aCSF). Sucrose aCSF contained (in mM): 185 sucrose, 25 NaHCO_3_, 2.5 KCl, 25 glucose, 1.25 NaH_2_PO_4_, 10 MgCl_2_, 0.5 CaCl_2_; pH 7.4, 330 mOsm. After hemisecting the brain, both hemispheres were glued on a platina. Acute hippocampal slices were prepared on a VT1000 S or VT1200 S Vibratome (Leica, Germany). Acute slices were then transferred to a heated (32°C) and oxygenated artificial cerebrospinal fluid (normal aCSF) that contained (in mM): 125 NaCl, 25 NaHCO_3_, 2.5 KCl, 10 glucose, 2 CaCl_2_, 2 MgCl_2_; pH 7.4, 300 mOsm. Slices were incubated at 32°C for 30 minutes, following which the water bath was turned off and the slices were left to recover for an additional 30 minutes before beginning experiments. Slices were then maintained at room temperature for the rest of the day and slices were used for up to 6 hours following preparation.

### Electrophysiological recordings

Acute hippocampal slices were transferred to a recording chamber and held under a harp. The recording chamber was continuously perfused (2 mL/min) with oxygenated aCSF at room temperature (20 ± 2°C, mean ± SD). An upright microscope (BX50WI or BX61WI, Olympus) equipped with a 40X water-immersion objective was used to visualize the hippocampus. Whole-cell patch clamp recordings were performed from visually identified interneurons expressing TdTomato (Figs. 1 and 2), or from putative pyramidal, fast-spiking, and regular-spiking interneurons that were then functionally identified (Figs. 3 and 4). Recording electrodes were obtained from borosilicate glass filaments (TW150-4, World Precision Instruments) pulled on a P-97 Micropipette Puller (Sutter Instruments). Electrodes had resistance of 3 – 6 MΩ. These electrodes were filled with a solution composed of (in mM): 130 K-gluconate, 10 HEPES, 2 MgCl_2_.6H_2_O, 2 Mg_2_ATP, 0.3 NaGTP, 7 Na_2_-Phosphocreatine, 0.6 EGTA, 5 KCl; pH 7.2 and 295 mOsm. The liquid junction potential was not corrected. The electrophysiological signal was amplified with an Axopatch 200B or a MultiClamp 700B and digitized at 10 kHz with a Digidata 1322A (Axon Instruments). The data was recorded on personal computers equipped with Clampex 8.2 and 9.2 programs. The data was saved on a personal computer. Optogenetic stimulation was delivered through an optical fiber positioned in *stratum oriens* with a micromanipulator. Blue light (470 nm) was generated by a light-emitting diode (LED) and precisely delivered by a TTL signal originating from the digitizer and sent to the LED controller (WT&T Inc.).

### Biocytin revelation and confocal microscopy

Following whole-cell recordings, acute hippocampal slices were fixed with freshly prepared PBS solution containing 4% PFA and left in the fridge overnight. The fixed acute hippocampal slices were processed for biocytin revelation. Briefly, slices were rinsed with PBS (4 x 5 min), treated with H_2_O_2_ (0.3%, 30 min), permeabilized with Triton (1%, 1 hour) and exposed to a streptavidin-conjugated Alexa-633 (1:200, overnight). Slices were rinsed with PBS (4 x 5 min) and mounted on microscope slides with ProLong Gold (ThermoFisher Scientific). Slices were kept in the fridge for at least two weeks before confocal imaging. Microscope slides with recovered neurons were imaged under an upright confocal microscope (Axo Imager.Z2, Zeiss). The soma location was identified under a low magnification objective (5X). A 40X oil-immersion objective was used for image acquisition. Z-stacks were acquired through the full Z-axis, in a concentric manner from the soma. We followed axonal and dendritic branches to their termination zones.

### Analysis of single cell transcriptomic dataset

We used the single cell transcriptomic data set from Harris et al., 2018, accessed at: https://figshare.com/articles/dataset/Transcriptomic_analysis_of_CA1_inhibitory_interneurons/6 <u>198656</u>. Genes with no expression were eliminated, and we first focused on the genes determined to define interneurons subclasses. For each gene pair, the product of the expression level was computed. A filter of 50 – 400 neurons was set for putative cluster identification. The neurons with an expression product > 1 for individual gene pairs were then identified. The interquartile range and standard distance were measured from the X-Y coordinates of these neurons on the Figure 2 presented in Harris et al., (2018). We then ranked these putative subclusters based on the weighted average between the standard distance and the interquartile range to identify the top 50 gene pairs for each gene defining interneuron subclasses. While multiple genes and pairs of genes could in theory allow us to target the same clusters, we preferentially used those for which transgenic animals were already available. Despite not fitting the above criteria completely, we generated the *Sst;;Nos1* animals with prior knowledge that these animals identify a very scarce subtype of INs in the cortex, and likely with a low density in the hippocampus (28), hinting that this intersection might target a relatively sparse and well defined population of interneurons.

### Neurolucida reconstructions and anatomical analysis

Confocal images were used to reconstruct the morphology of biocytin-filled neurons with the Neurolucida 360 software. Following complete tracing of the neurites, 10 um thick contours were drawn over the entirety of the neuron. The border between *strata pyramidale* and *radiatum* was used as a landmark to measure perpendicular distances. Axonal density was then quantified by Neurolucida Explorer, which calculated the total axon length in each contour. These lengths were averaged across all cells for *Sst*-INs, *Sst;;Tac1*-INs, *Ndnf;;Nkx2-1*-INs, *Chrna2*-INs and *Sst;;Nos1*-INs. To calculate the cumulative distribution of axon length for each cell type, the total length of axon in each contour was normalized to the summed axon length for that cell. These normalized length distributions were then averaged across multiple cells for individual genotypes.

### Data analysis, statistical tests and k-means analysis

Electrophysiological data was analyzed in Clampfit 10.3 (Molecular Devices) and results were compiled in Microsoft Excel. Kolmogorov–Smirnov tests on anatomical and electrophysiological parameters were performed in GraphPad Prism for macOS (Version 9.5.1). P-values reported in Supplementary Tables 1 – 2 were corrected for multiple comparison using the Holm-Bonferroni method. For normally distributed data, Student’s t-test was used to evaluate statistical significance. For non-normally distributed data, a Mann-Whitney test was used. Principal component analysis (PCA) was carried out for 81 neurons using the following 8 electrophysiological properties: action potential amplitude, threshold, maximum rate of decay, maximum rate of rise, full width at half maximum, afterhyperpolarization maximal amplitude, sag amplitude and rebound depolarization. Scikit’s sklearn.decomposition.PCA function was used to calculate the transformation of this dataset. The absolute values in the eigenvectors corresponding to each property were used to determine the importance of the features within each principal component (Supplementary Table 3). The first four principal components accounted for more than 90% of the variance of the dataset and so were used for subsequent K-means clustering analysis. For K-means clustering, scikit’s sklearn.cluster.kmeans function was firstly used to determine the optimal value of k via the elbow method. Scipy’s scipy.cluster.vq.kmeans2 function was used to distribute the dataset into 2 clusters using the k-means algorithm. The algorithm is optimized to form clusters with minimal Euclidean distance between each data point and its assigned centroid, which represents the arithmetic mean of the data points in a particular cluster.

## Acknowledgements

We thank Dr. Bernardo Rudy for providing the *Ndnf*-Flp mouse, and Raquel Moya for help with PCA and k-means analyses. SC was supported by a senior biomedical postdoctoral fellowship from the Charles H. Revson Foundation, a postdoctoral fellowship from the Fonds de Recherche en Santé Québec, a K99/R00 Pathway to Independence Award from NIMH (1K99MH126157-01), and the Andrew Ellis and Emily Segal Investigator Grant from the Brain and Behavior Research Foundation. RM was supported by the NINDS (P01NS074972). KK received grants from Swedish Research Council (2022-01245), Swedish Brain Foundation (FO2022 - 0018) and Olle Engkvist Foundation (462193024). RWT received grants from the NINDS (1U19NS107616-02), NIDA (R01 DA040484-04) and NIMH (R01 MH071739-15). We thank the New York University Langone Medical Center Rodent Genetic Engineering Laboratory (P30CA016087).

---

**Supplementary Figure 1:**
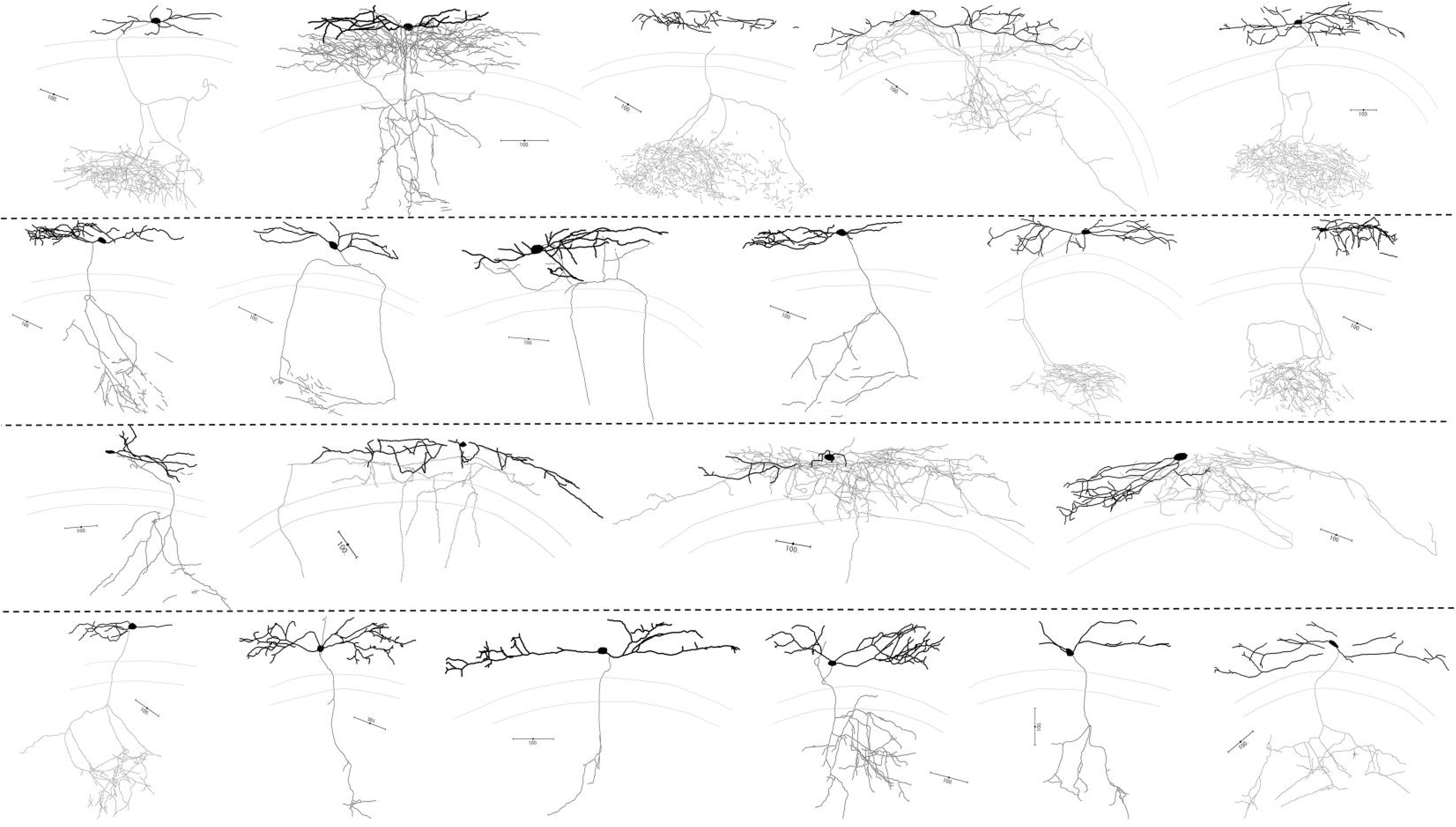
Neurolucida reconstructions of *Sst*-Ins. Neurolucida reconstructions of representative biocytin-filled *Sst*-INs. Whole cell patch clamp recorded soma in O/A. Axon is shown in gray, and the dendrites are shown in black. All scale bars represent 100 μm.

**Supplementary Figures 2:**
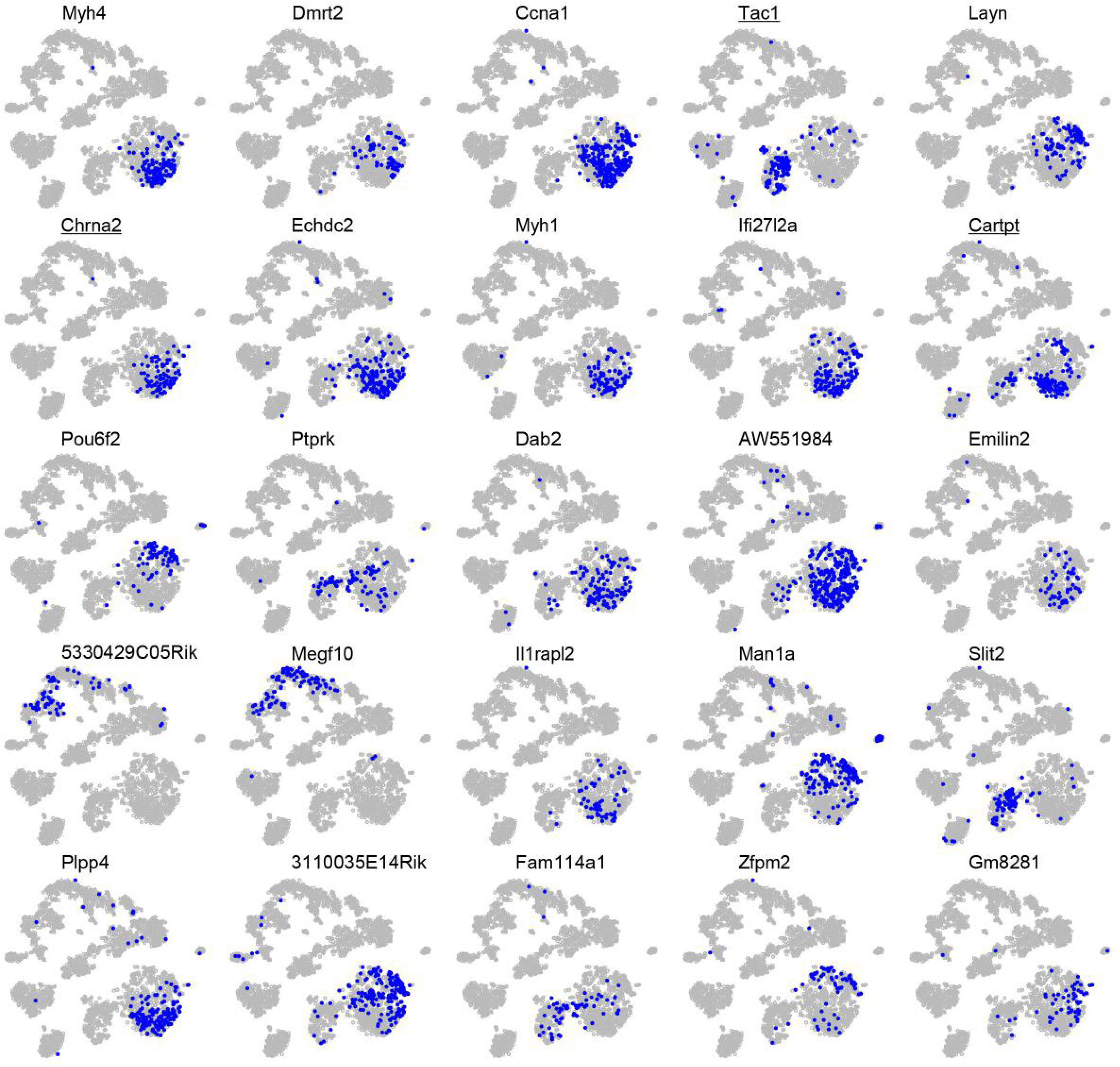
Spatial distribution analysis of the transcriptomic dataset. Top 25 hits minimizing for spatial dispersion for pairs of genes (shown above maps) combined with *Sst*. Multiple genes fit the criteria and could in principle be used. Underlined genes identify genes for which transgenic mouse models exist.

**Supplementary Figure 3:**
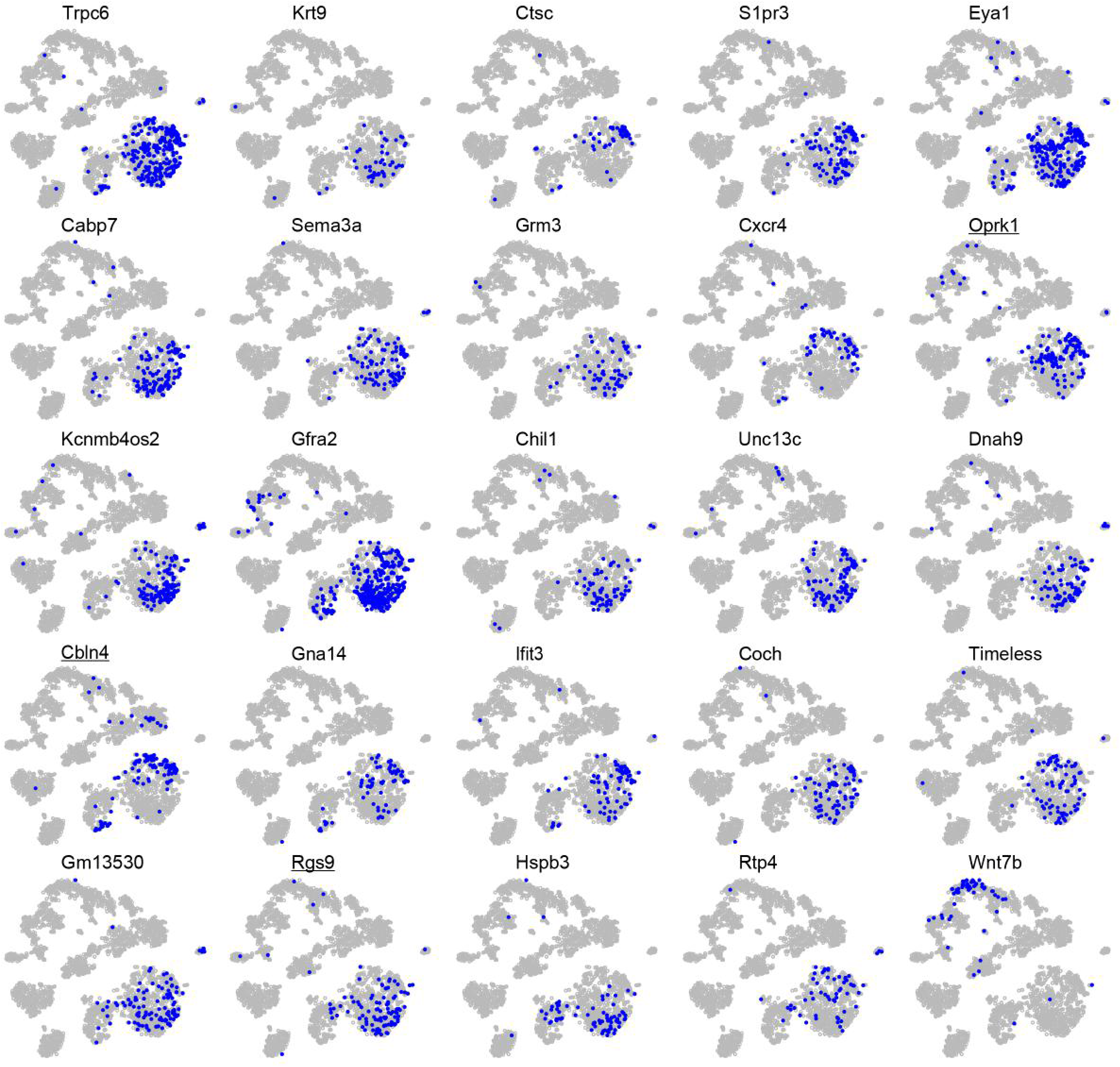
Spatial distribution analysis of the transcriptomic dataset, continued. Next 25 hits in the spatial distribution analysis shown in Fig. S2.

**Supplementary Figure 4:**
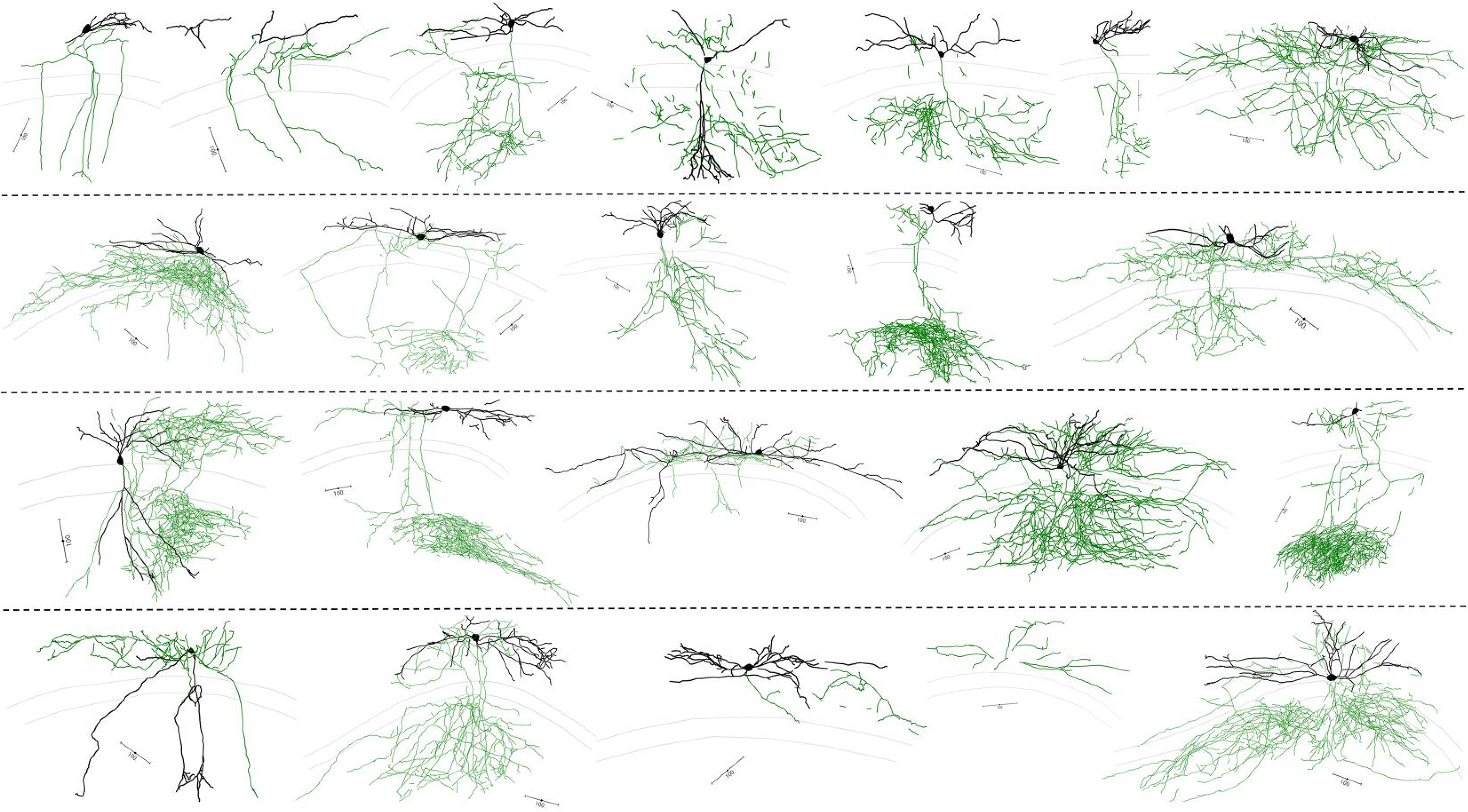
Neurolucida reconstructions of *Sst;;Tac1*-Ins. Neurolucida reconstructions of biocytin-filled *Sst;;Tac1*-INs. Axon is shown in green, and the dendrites are shown in black. All scale bars represent 100 μm.

**Supplementary Figure 5:**
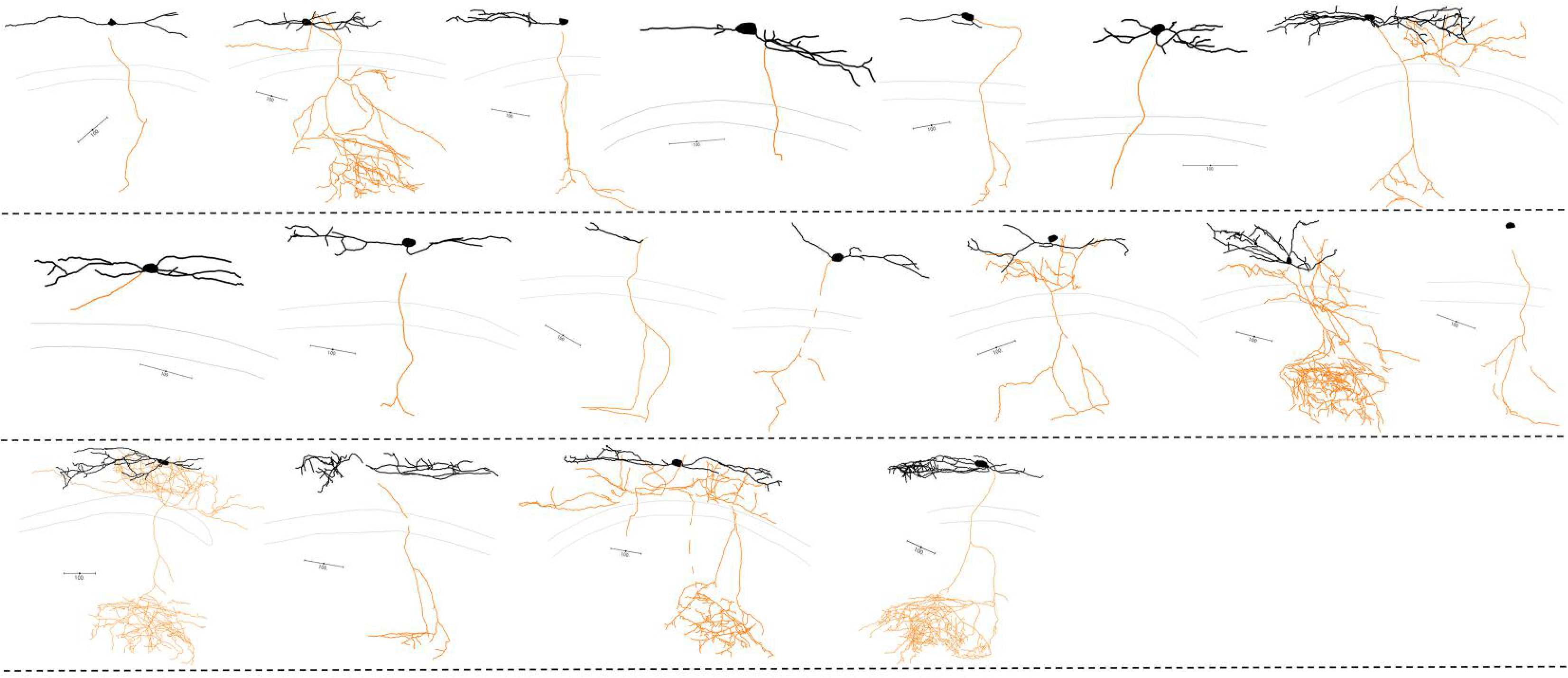
Neurolucida reconstructions of *Ndnf;;Nkx2-1*-Ins. Neurolucida reconstructions of biocytin-filled *Ndnf;;Nkx2-1*-INs. Axon is shown in orange, and the dendrites are shown in black. All scale bars represent 100 μm.

**Supplementary Figure 6:**
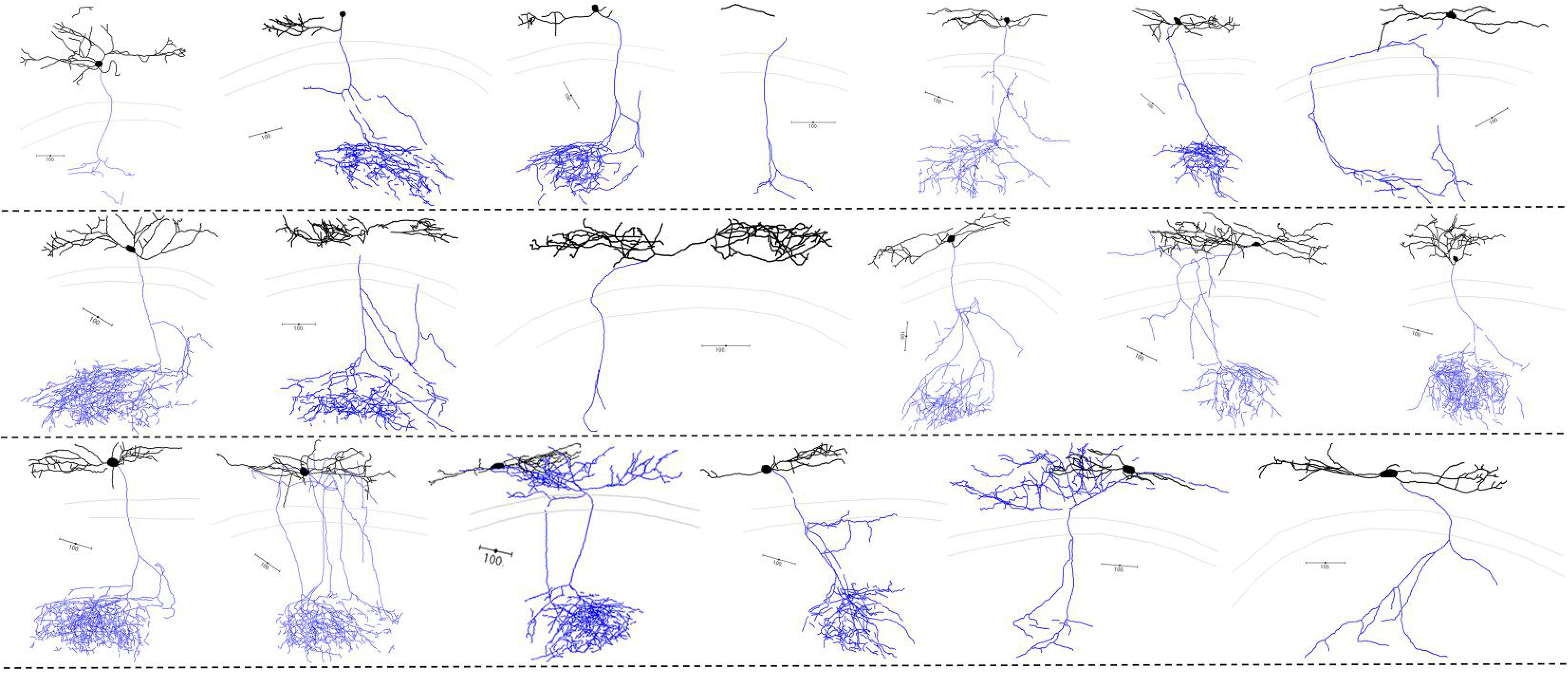
Neurolucida reconstructions of *Chrna2*-Ins. Neurolucida reconstructions of biocytin-filled *Chrna2*-INs. Axon is shown in blue, and the dendrites are shown in black. All scale bars represent 100 μm.

**Supplementary Figure 7:**
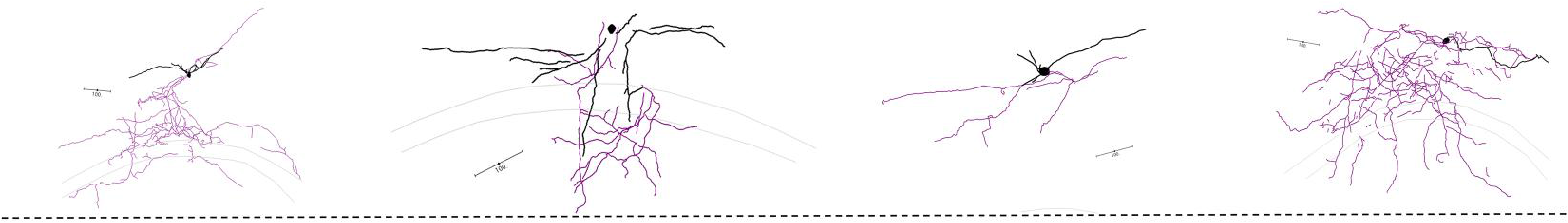
Neurolucida reconstructions of *Sst;;Nos1*-Ins. Neurolucida reconstructions of biocytin-filled *Sst;;Nos1*-INs. Axon is shown in purple, and the dendrites are shown in black. All scale bars represent 100 μm.

**Supplementary Figure 8:**
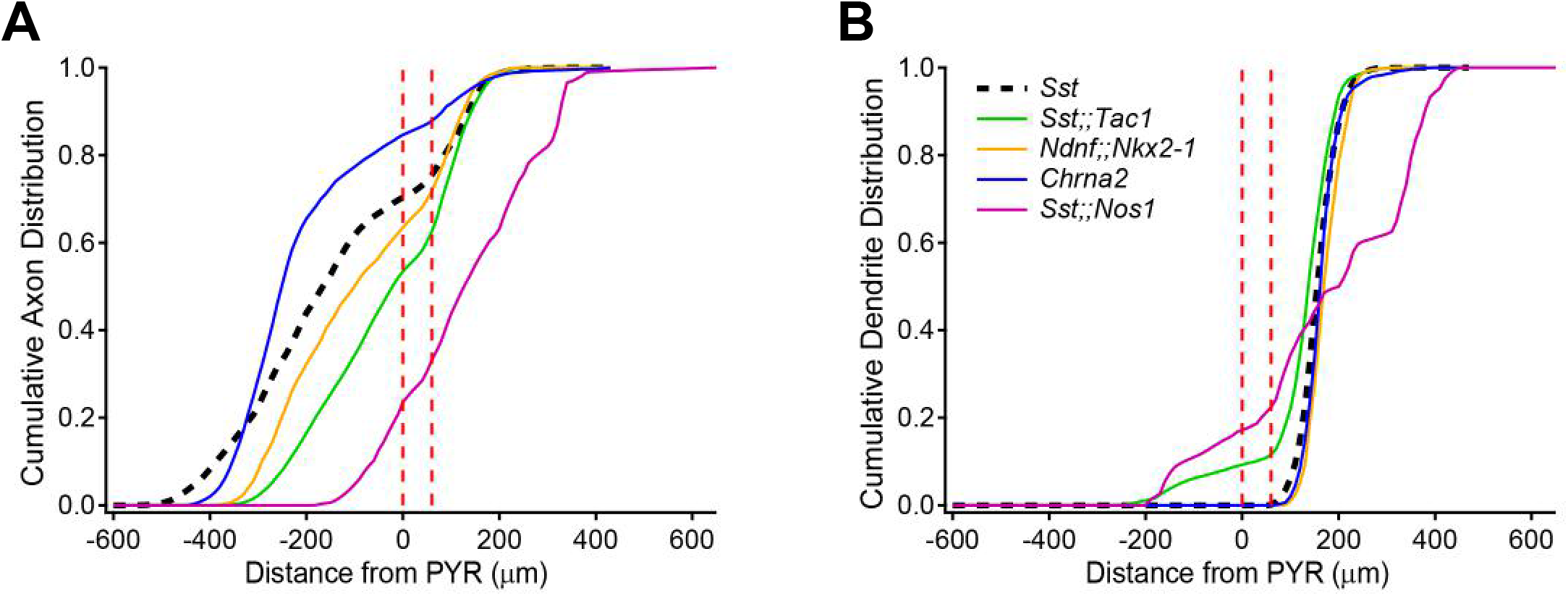
Axonal and dendritic distributions. **A**, Cumulative axonal distribution for all neurons recorded, alternative representation to Fig. 1H. **B**, Cumulative dendritic distribution for all neurons recorded.

**Supplementary Figure 9:**
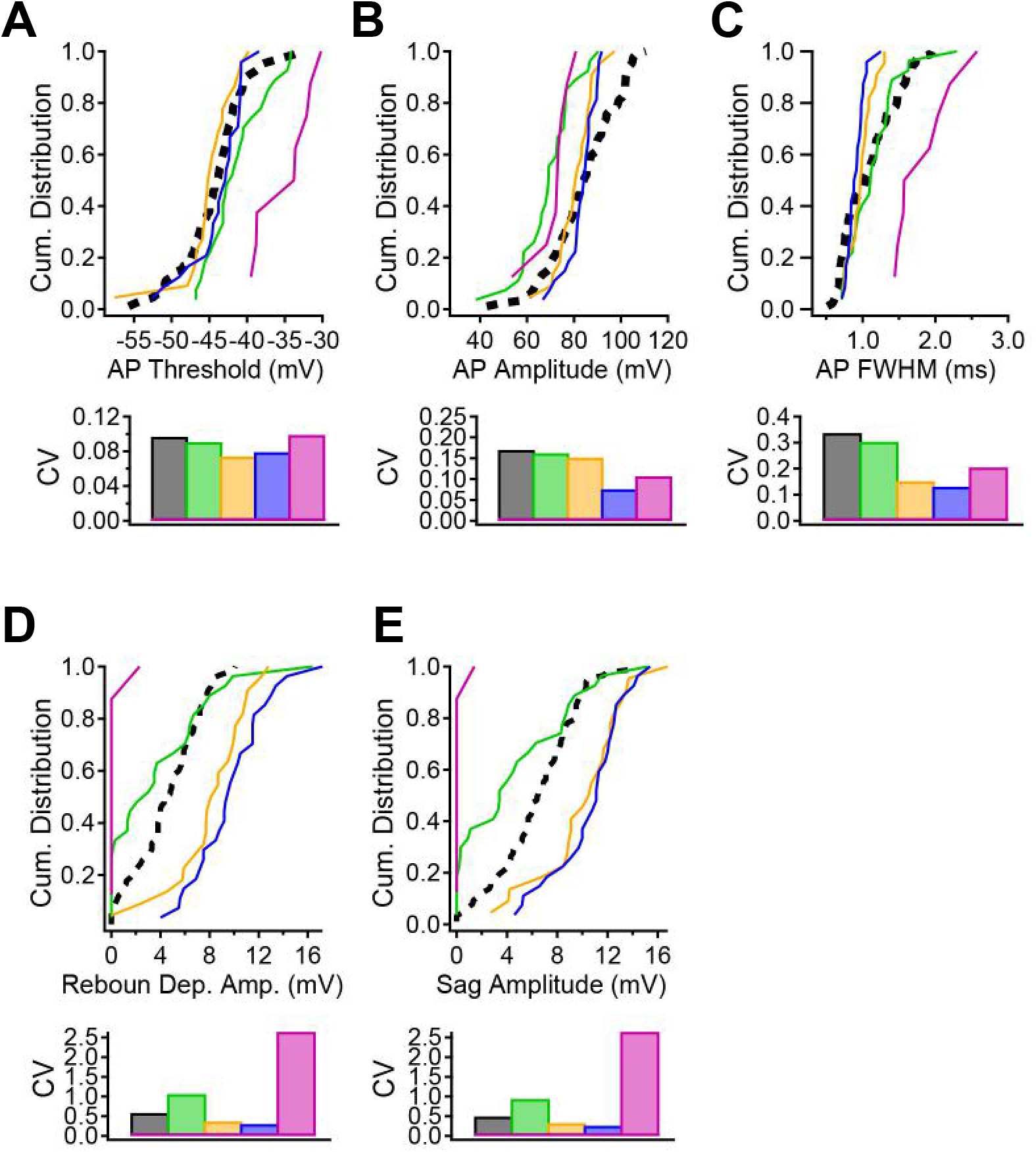
Analysis of electrophysiological parameters used for clustering. **A – E**, Cumulative distributions of AP threshold (A), AP amplitude (B), AP full width at half maximum (C), rebound depolarization amplitude (D) and sag amplitude (E) for all neurons recorded in this study. The coefficient of variation measured across all neurons is shown below each graph. The combination of these 5 parameters and the 3 parameters reported in Figure 2 were used for the cluster analysis.

**Supplementary Table 1:** p-values for statistical comparisons of anatomical parameters. P-values reported for Kolmogorov-Smirnov tests followed by Holm-Bonferroni correction.

| **p-values** |  | ***Sst*** | ***Sst;;Tac1*** | ***Ndnf;;Nkx2-1*** | ***Chrna2*** |
| --- | --- | --- | --- | --- | --- |
| **axons** | ***Sst*** |  |  |  |  |
|  | ***Sst;;Tac1*** | 0.3417 |  |  |  |
|  | ***Ndnf;;Nkx2*** | 0.3417 | 0.3417 |  |  |
|  | ***Chrna2*** | 0.0036 | 0.005 | 0.0064 |  |
|  | ***Sst;;Nos1*** | 0.000048 | 0.000147 | 0.00001 | 0.00001 |
| **dendrites** | ***Sst*** |  |  |  |  |
|  | ***Sst;;Tac1*** | 0.0032 |  |  |  |
|  | ***Ndnf;;Nkx2*** | 1 | 0.00068 |  |  |
|  | ***Chrna2*** | 1 | 0.0018 | 1 |  |
|  | ***Sst;;Nos1*** | 0.0305 | 0.2416 | 0.0112 | 0.015 |

**Supplementary Table 2:** p-values for statistical comparisons of electrophysiological parameters. P-values reported for Kolmogorov-Smirnov tests followed by Holm-Bonferroni correction.

| p-values |  | *Sst* | *Sst;;Tac1* | *Ndnf;;Nkx2-1* | *Chrna2* |
| --- | --- | --- | --- | --- | --- |
| ahp_max | *Sst* |  |  |  |  |
|  | *Sst;;Tac1* | 1 |  |  |  |
|  | *Ndnf;;Nkx2* | 0.2709 | 1 |  |  |
|  | *Chrna2* | 0.4568 | 0.5976 | 0.051 |  |
|  | *Sst;;Nos1* | 1 | 1 | 0.567 | 1 |
| ap_amp | *Sst* |  |  |  |  |
|  | *Sst;;Tac1* | 0.00005 |  |  |  |
|  | *Ndnf;;Nkx2* | 0.21 | 0.014 |  |  |
|  | *Chrna2* | 0.18 | 0.00003 | 0.614 |  |
|  | *Sst;;Nos1* | 0.093 | 0.614 | 0.21 | 0.009 |
| ap_thresh | *Sst* |  |  |  |  |
|  | *Sst;;Tac1* | 0.2855 |  |  |  |
|  | *Ndnf;;Nkx2* | 0.8342 | 0.0486 |  |  |
|  | *Chrna2* | 0.8342 | 0.4924 | 0.4924 |  |
|  | *Sst;;Nos1* | 0.0001 | 0.0308 | 0.000144 | 0.000264 |
| ap_fwhm | *Sst* |  |  |  |  |
|  | *Sst;;Tac1* | 0.5658 |  |  |  |
|  | *Ndnf;;Nkx2* | 0.3138 | 0.1336 |  |  |
|  | *Chrna2* | 0.0075 | 0.0054 | 0.5658 |  |
|  | *Sst;;Nos1* | 0.0021 | 0.0008 | 0.000144 | 0.00012 |
| sag | *Sst* |  |  |  |  |
|  | *Sst;;Tac1* | 0.037 |  |  |  |
|  | *Ndnf;;Nkx2* | 0.000252 | 0.0016 |  |  |
|  | *Chrna2* | 0.00001 | 0.000252 | 0.9484 |  |
|  | *Sst;;Nos1* | 0.000088 | 0.0168 | 0.000112 | 0.000081 |
| ap_maxdecay | *Sst* |  |  |  |  |
|  | *Sst;;Tac1* | 0.0072 |  |  |  |
|  | *Ndnf;;Nkx2* | 0.0344 | 0.0135 |  |  |
|  | *Chrna2* | 0.0028 | 0.000405 | 0.0344 |  |
|  | *Sst;;Nos1* | 0.0042 | 0.0055 | 0.000405 | 0.00009 |
| ap_maxrise | *Sst* |  |  |  |  |
|  | *Sst;;Tac1* | 0.003 |  |  |  |
|  | *Ndnf;;Nkx2* | 0.4408 | 0.0016 |  |  |
|  | *Chrna2* | 0.0884 | 0.00003 | 0.0884 |  |
|  | *Sst;;Nos1* | 0.006 | 0.349 | 0.0021 | 0.000189 |
| rebound dep | *Sst* |  |  |  |  |
|  | *Sst;;Tac1* | 0.1044 |  |  |  |
|  | *Ndnf;;Nkx2* | 0.000096 | 0.0052 |  |  |
|  | *Chrna2* | 0.00001 | 0.000096 | 0.3972 |  |
|  | *Sst;;Nos1* | 0.00052 | 0.0558 | 0.000276 | 0.000081 |

**Supplementary Table 3:** Contribution of individual parameter to the principal components.

|  | PC1_feat | PC2_feat | PC3_feat | PC4_feat | PC5_feat | PC6_feat | PC7_feat | PC8_feat |
| --- | --- | --- | --- | --- | --- | --- | --- | --- |
| ap_thresh | 0.38 | 0.39 | 0.35 | 0.44 | 0.38 | 0.35 | 0.35 | 0.05 |
| ap_amp | 0.09 | 0.17 | 0.38 | 0.14 | 0.44 | 0.33 | 0.32 | 0.63 |
| ap_fwhm | 0.54 | 0.13 | 0.02 | 0.03 | 0.09 | 0.48 | 0.48 | 0.47 |
| ap_maxrise | 0.12 | 0.68 | 0.56 | 0.22 | 0.02 | 0.24 | 0.04 | 0.32 |
| ap_maxdecay | 0.66 | 0.01 | 0.11 | 0.53 | 0.08 | 0.01 | 0.24 | 0.45 |
| sag | 0.17 | 0.08 | 0.26 | 0.08 | 0.06 | 0.68 | 0.64 | 0.14 |
| rebound dep | 0.18 | 0.57 | 0.55 | 0.42 | 0.28 | 0.14 | 0.27 | 0.02 |
| ahp_max | 0.22 | 0.12 | 0.20 | 0.52 | 0.75 | 0.07 | 0.04 | 0.23 |

## References

1. T. F. Freund, G. Buzsaki, Interneurons of the hippocampus. Hippocampus 6, 347–470 (1996).

2. K. A. Pelkey et al., Hippocampal GABAergic Inhibitory Interneurons. Physiol Rev 97, 1619–1747 (2017).

3. K. D. Harris et al., Classes and continua of hippocampal CA1 inhibitory neurons revealed by single-cell transcriptomics. PLoS Biol 16, e2006387 (2018).

4. B. Rudy, G. Fishell, S. Lee, J. Hjerling-Leffler, Three groups of interneurons account for nearly 100% of neocortical GABAergic neurons. Dev Neurobiol 71, 45–61 (2011).

5. T. Klausberger, P. Somogyi, Neuronal diversity and temporal dynamics: the unity of hippocampal circuit operations. Science 321, 53–57 (2008).

6. S. Royer et al., Control of timing, rate and bursts of hippocampal place cells by dendritic and somatic inhibition. Nat Neurosci 15, 769–775 (2012).

7. F. Pouille, M. Scanziani, Routing of spike series by dynamic circuits in the hippocampus. Nature 429, 717–723 (2004).

8. J. C. Lacaille, A. L. Mueller, D. D. Kunkel, P. A. Schwartzkroin, Local circuit interactions between oriens/alveus interneurons and CA1 pyramidal cells in hippocampal slices: electrophysiology and morphology. J Neurosci 7, 1979–1993 (1987).

9. M. Lovett-Barron et al., Regulation of neuronal input transformations by tunable dendritic inhibition. Nat Neurosci 15, 423–430, S421-423 (2012).

10. A. D. Milstein et al., Inhibitory Gating of Input Comparison in the CA1 Microcircuit. Neuron 87, 1274–1289 (2015).

11. R. N. Leao et al., OLM interneurons differentially modulate CA3 and entorhinal inputs to hippocampal CA1 neurons. Nat Neurosci 15, 1524–1530 (2012).

12. S. Siwani et al., OLMalpha2 Cells Bidirectionally Modulate Learning. Neuron 99, 404–412 e403 (2018).

13. C. Muller, S. Remy, Dendritic inhibition mediated by O-LM and bistratified interneurons in the hippocampus. Front Synaptic Neurosci 6, 23 (2014).

14. S. A. Booker, I. Vida, Morphological diversity and connectivity of hippocampal interneurons. Cell Tissue Res 373, 619–641 (2018).

15. L. Katona et al., Sleep and movement differentiates actions of two types of somatostatin-expressing GABAergic interneuron in rat hippocampus. Neuron 82, 872–886 (2014).

16. R. Chittajallu et al., Dual origins of functionally distinct O-LM interneurons revealed by differential 5-HT(3A)R expression. Nat Neurosci 16, 1598–1607 (2013).

17. S. Mikulovic, C. E. Restrepo, M. M. Hilscher, K. Kullander, R. N. Leao, Novel markers for OLM interneurons in the hippocampus. Front Cell Neurosci 9, 201 (2015).

18. P. Somogyi, T. Klausberger, Defined types of cortical interneurone structure space and spike timing in the hippocampus. J Physiol 562, 9–26 (2005).

19. Y. Ma, H. Hu, A. S. Berrebi, P. H. Mathers, A. Agmon, Distinct subtypes of somatostatin-containing neocortical interneurons revealed in transgenic mice. J Neurosci 26, 5069–5082 (2006).

20. S. Chamberland et al., Brief synaptic inhibition persistently interrupts firing of fast-spiking interneurons. Neuron 10.1016/j.neuron.2023.01.017 (2023).

21. J. Artinian, J. C. Lacaille, Disinhibition in learning and memory circuits: New vistas for somatostatin interneurons and long-term synaptic plasticity. Brain Res Bull 141, 20–26 (2018).

22. I. Katona, L. Acsady, T. F. Freund, Postsynaptic targets of somatostatin-immunoreactive interneurons in the rat hippocampus. Neuroscience 88, 37–55 (1999).

23. H. Xu, H. Y. Jeong, R. Tremblay, B. Rudy, Neocortical somatostatin-expressing GABAergic interneurons disinhibit the thalamorecipient layer 4. Neuron 77, 155–167 (2013).

24. Z. Yao et al., A taxonomy of transcriptomic cell types across the isocortex and hippocampal formation. Cell 184, 3222–3241 e3226 (2021).

25. R. Tremblay, S. Lee, B. Rudy, GABAergic Interneurons in the Neocortex: From Cellular Properties to Circuits. Neuron 91, 260–292 (2016).

26. B. Tasic et al., Shared and distinct transcriptomic cell types across neocortical areas. Nature 563, 72–78 (2018).

27. L. Tricoire et al., A blueprint for the spatiotemporal origins of mouse hippocampal interneuron diversity. J Neurosci 31, 10948–10970 (2011).

28. M. He et al., Strategies and Tools for Combinatorial Targeting of GABAergic Neurons in Mouse Cerebral Cortex. Neuron 92, 555 (2016).

29. G. Maccaferri, C. J. McBain, The hyperpolarization-activated current (Ih) and its contribution to pacemaker activity in rat CA1 hippocampal stratum oriens-alveus interneurones. J Physiol 497 ( **Pt 1**), 119–130 (1996).

30. J. J. Tukker, P. Fuentealba, K. Hartwich, P. Somogyi, T. Klausberger, Cell type-specific tuning of hippocampal interneuron firing during gamma oscillations in vivo. J Neurosci 27, 8184–8189 (2007).

31. T. Klausberger et al., Spike timing of dendrite-targeting bistratified cells during hippocampal network oscillations in vivo. Nat Neurosci 7, 41–47 (2004).

32. C. K. Pfeffer, M. Xue, M. He, Z. J. Huang, M. Scanziani, Inhibition of inhibition in visual cortex: the logic of connections between molecularly distinct interneurons. Nat Neurosci 16, 1068–1076 (2013).

33. P. Parra, A. I. Gulyas, R. Miles, How many subtypes of inhibitory cells in the hippocampus? Neuron 20, 983–993 (1998).

34. E. H. Buhl, K. Halasy, P. Somogyi, Diverse sources of hippocampal unitary inhibitory postsynaptic potentials and the number of synaptic release sites. Nature 368, 823–828 (1994).

35. E. H. Buhl, T. Szilagyi, K. Halasy, P. Somogyi, Physiological properties of anatomically identified basket and bistratified cells in the CA1 area of the rat hippocampus in vitro. Hippocampus 6, 294–305 (1996).

36. G. F. Turi et al., Vasoactive Intestinal Polypeptide-Expressing Interneurons in the Hippocampus Support Goal-Oriented Spatial Learning. Neuron 101, 1150–1165 e1158 (2019).

37. S. Chamberland, C. Salesse, D. Topolnik, L. Topolnik, Synapse-specific inhibitory control of hippocampal feedback inhibitory circuit. Front Cell Neurosci 4, 130 (2010).

38. L. Acsady, T. J. Gorcs, T. F. Freund, Different populations of vasoactive intestinal polypeptide-immunoreactive interneurons are specialized to control pyramidal cells or interneurons in the hippocampus. Neuroscience 73, 317–334 (1996).

39. L. Tyan et al., Dendritic inhibition provided by interneuron-specific cells controls the firing rate and timing of the hippocampal feedback inhibitory circuitry. J Neurosci 34, 4534–4547 (2014).

40. J. Winterer et al., Single-cell RNA-Seq characterization of anatomically identified OLM interneurons in different transgenic mouse lines. Eur J Neurosci 50, 3750–3771 (2019).

41. A. Paul et al., Transcriptional Architecture of Synaptic Communication Delineates GABAergic Neuron Identity. Cell 171, 522–539 e520 (2017).

42. C. Foldy et al., Single-cell RNAseq reveals cell adhesion molecule profiles in electrophysiologically defined neurons. Proc Natl Acad Sci U S A 113, E5222–5231 (2016).

43. A. Kepecs, G. Fishell, Interneuron cell types are fit to function. Nature 505, 318–326 (2014).

44. E. S. Boyden, F. Zhang, E. Bamberg, G. Nagel, K. Deisseroth, Millisecond-timescale, genetically targeted optical control of neural activity. Nat Neurosci 8, 1263–1268 (2005).

45. Z. J. Huang, W. Yu, C. Lovett, S. Tonegawa, Cre/loxP recombination-activated neuronal markers in mouse neocortex and hippocampus. Genesis 32, 209–217 (2002).

46. T. L. Daigle et al., A Suite of Transgenic Driver and Reporter Mouse Lines with Enhanced Brain-Cell-Type Targeting and Functionality. Cell 174, 465–480 e422 (2018).

47. H. Taniguchi et al., A resource of Cre driver lines for genetic targeting of GABAergic neurons in cerebral cortex. Neuron 71, 995–1013 (2011).

48. L. E. Fenno et al., Comprehensive Dual- and Triple-Feature Intersectional Single-Vector Delivery of Diverse Functional Payloads to Cells of Behaving Mammals. Neuron 107, 836–853 e811 (2020).

49. V. S. Sohal, F. Zhang, O. Yizhar, K. Deisseroth, Parvalbumin neurons and gamma rhythms enhance cortical circuit performance. Nature 459, 698–702 (2009).

50. J. Dimidschstein et al., A viral strategy for targeting and manipulating interneurons across vertebrate species. Nat Neurosci 19, 1743–1749 (2016).

51. G. Pouchelon et al., A versatile viral toolkit for functional discovery in the nervous system. Cell Rep Methods 2, 100225 (2022).

52. D. Vormstein-Schneider et al., Viral manipulation of functionally distinct interneurons in mice, non-human primates and humans. Nat Neurosci 23, 1629–1636 (2020).

53. Y. Qian et al., Programmable RNA sensing for cell monitoring and manipulation. Nature 610, 713–721 (2022).

54. S. J. Wu et al., Cortical somatostatin interneuron subtypes form cell-type specific circuits. bioRxiv 10.1101/2022.09.29.510081, 2022.2009.2029.510081 (2022).

55. R. E. Hostetler, H. Hu, A. Agmon, Genetically Defined Subtypes of Layer 5 Somatostatin-Containing Cortical Interneurons. bioRxiv 10.1101/2023.02.02.526850, 2023.2002.2002.526850 (2023).

56. J. A. Harris et al., Anatomical characterization of Cre driver mice for neural circuit mapping and manipulation. Front Neural Circuits 8, 76 (2014).

57. Q. Xu, M. Tam, S. A. Anderson, Fate mapping Nkx2.1-lineage cells in the mouse telencephalon. J Comp Neurol 506, 16–29 (2008).

58. L. Madisen et al., A robust and high-throughput Cre reporting and characterization system for the whole mouse brain. Nat Neurosci 13, 133–140 (2010).

59. L. Madisen et al., Transgenic mice for intersectional targeting of neural sensors and effectors with high specificity and performance. Neuron 85, 942–958 (2015).

60. L. Madisen et al., A toolbox of Cre-dependent optogenetic transgenic mice for light-induced activation and silencing. Nat Neurosci 15, 793–802 (2012).

